# Restoring beneficial ancestral genomic sites enhances broad disease resistance in rice

**DOI:** 10.64898/2026.11.22.768253

**Authors:** Jiyang Wang, Shuo Liu, Yuange Duan, Tianyi Li, Huarong Liu, Dongyu Shi, Baohui Mou, Jun Yang, Danbing Shan, Xinhang Zheng, Guosheng Zhao, Shanzhi Wang, Lin Zhu, Qiming Wu, Menghan Qu, Chunyu Yang, Mei Zhao, Xuankun Li, Shanlin Liu, Fang Xue, Shiyong Zhang, Hai Wang, Sheng Yang He, Wenxian Sun

## Abstract

Crop improvement has long been constrained to genetic variations within cultivated germplasms and their wild relatives, which capture only a small fraction of historical protein diversity accumulated over evolutionary processes. Here, we propose the “Beneficial Ancestral Genomic Sites” concept, hypothesizing that restoring conserved ancestral acidic residues at phosphorylatable sites in plant proteins could substantially improve plant performance. We experimentally validate this concept with key immune-associated NADPH oxidases OsRbohB and OsRbohI to elevate broad-spectrum disease resistance in rice. Field-level testing of two competing models for growth-defense tradeoffs in plants strongly supports mitigating tradeoffs in modern agricultural settings. Our work unlocks a hidden, untapped reservoir of elite molecular variations inaccessible from conventional breeding resources, establishing a potentially transformative strategy for next-generation crop improvement under modern agriculture contexts.

## Introduction

Crop improvement has traditionally relied on natural variation in cultivated germplasms and wild relatives^1,2^. However, the full evolutionary history of a species encompasses a far broader spectrum of molecular and phenotypic states than is represented by currently available breeding materials. Most historical variants that arose in ancestral or allied lineages, including allelic states that were once adaptive in ancient ecological contexts, have been lost from these modern breeding resources because of genetic drift, demographic bottlenecks, linked selection or long-term shifts in environmental selection pressures^3–5^. As a result, the standing genetic variation available to plant breeders represents only a narrow, time-sampled subset of the historical variation explored during species evolution. Genetic variants that are absent from existing populations cannot be recovered through screening of contemporary genetic diversity and are unlikely to arise de novo within the typical timescales of crop improvement^6,7^. This lost historical repertoire therefore represents a largely untapped source of potentially useful variation, but strategies to infer, reconstruct and test such states for crop improvement remain limited.

Some historical genetic variation can be inferred from extant protein sequences^8,9^. A subset of existing phosphorylation sites on serine, threonine and tyrosine evolved from ancestral aspartate or glutamate residues^10^. In this evolutionary process, acidic residues that usually provide a stable negative charge under physiological conditions were replaced by phosphorylatable residues, converting a constitutive charge state into a regulatory state in which negative charge is dynamically introduced by post-translational phosphorylation^10^. These acidic-to-phosphorylatable residues (hereafter referred to as candidate evolutionarily acquired phosphorylation sites or EAPSs) establish a striking evolutionary contrast between the constitutively charged ancestral acidic state and the derived phosphorylation-dependent regulatory state. Whether this evolutionary divergence harbors recoverable historical variation with biological and agronomic value remains experimentally untested.

Disease resistance provides a uniquely stringent biological system to test this hypothesis. Immune regulators are shaped by fundamentally opposing demands: defense responses must be activated rapidly enough to constrain pathogen infection, yet remain tightly restrained to minimize fitness costs associated with growth and reproduction^11,12^. These counteracting selective pressures can precisely tune the amplitude, timing and activation threshold of immune signaling, making EAPSs a potential molecular readout of historical regulatory rewiring in plant immunity. Grasses, including major cereal crops, have a complex evolutionary and domestication history marked by deep phylogenetic divergence, independent crop domestication events and selection pressures from diverse pathogens^13–15^. Despite these diverse evolutionary trajectories, the prevalence of derived phosphoregulatory states in grasses and the potential of ancestral residues to reveal disease resistance-associated variation have not been systematically explored.

## Results

### Candidate evolutionarily acquired phosphorylation sites are widespread in grasses

To determine whether acidic-to-phosphorylatable residue substitutions are pervasive in grass protein evolution (Figure S1), we first identified orthologous proteins across 17 representative plant species spanning gymnosperms, basal angiosperms, eudicots and monocots, including five Poaceae species for subsequent grass-lineage comparisons (Figure 1A and Figure S2). This analysis yielded 15,804 orthologous groups (OGs) containing at least two proteins, and the protein sequences within each OG were aligned for residue-level comparison (Table S1).

**Figure 1.**
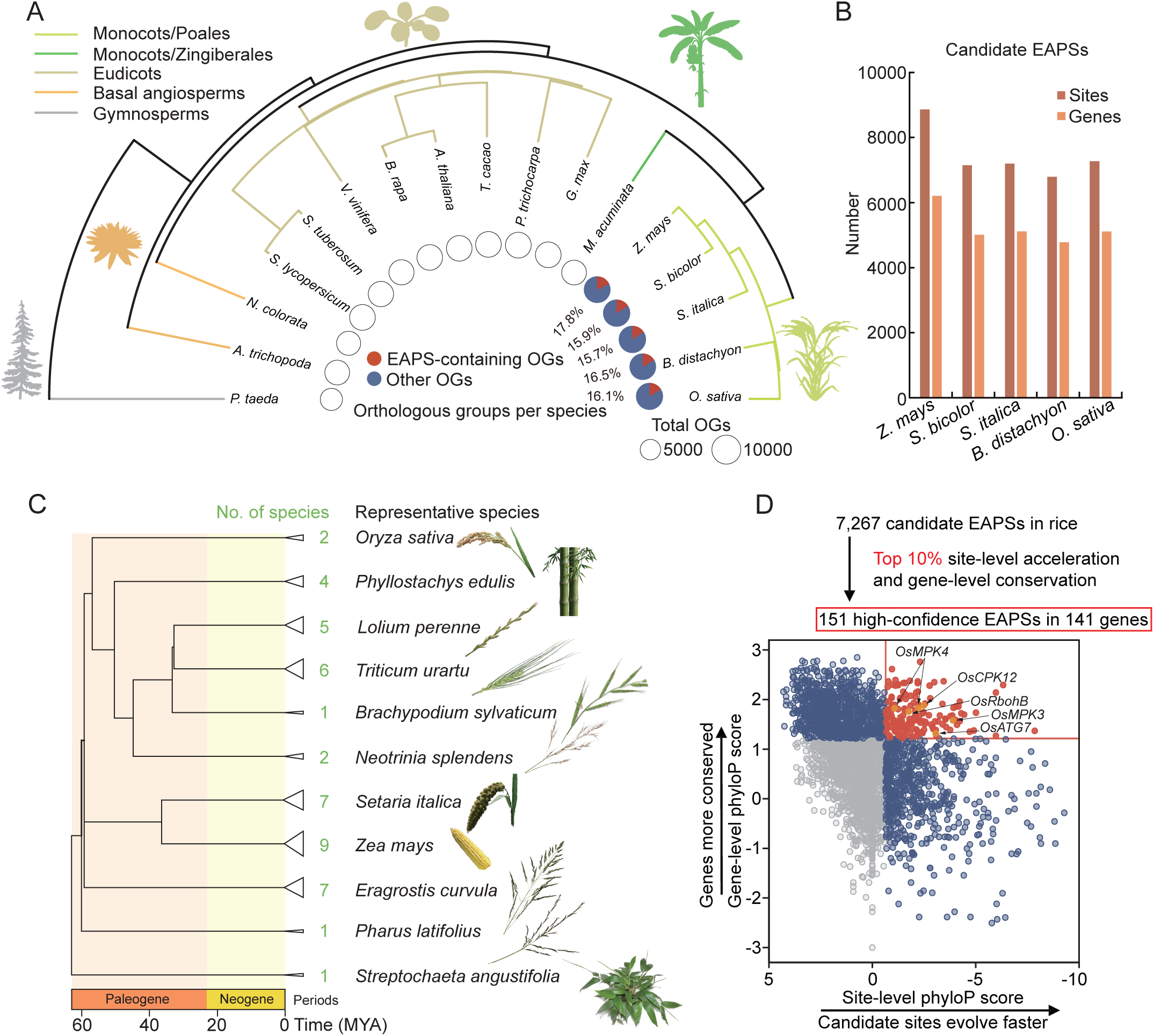
Evolutionary analysis reveals widespread EAPSs in grasses and prioritizes candidate sites in rice. (A) Phylogenetic framework of 17 representative seed plant species to identify candidate evolutionarily acquired phosphorylation sites (EAPSs). Branch colours indicate major lineages: gymnosperms, early-diverging angiosperms, eudicots, non-Poales monocots and Poales. Candidate EAPSs were defined as orthologous positions at which an inferred ancestral acidic residue, Asp or Glu, was replaced by a phosphorylatable residue, Ser, Thr or Tyr, in at least one Poaceae species. Pie charts show the proportion of orthologous groups (OGs) containing at least one candidate EAPS in each Poaceae species; orange sectors indicate EAPS-containing OGs and blue sectors indicate other OGs. Circle size indicates the total number of OGs analysed in each species. (B) Numbers of candidate EAPSs and corresponding genes identified in five representative Poaceae species. Sites indicate acidic-to-phosphorylatable residue substitutions and genes indicate protein-coding genes containing at least one candidate EAPS. (C) Phylogenetic tree of the 45 Poaceae species for whole-genome alignment and phyloP-based prioritization of candidate rice EAPSs. Each terminal branch represents lineage collapsed by species with divergence times within 30 MYA, with green numbers indicating the number of species represented. The bottom color bar denotes geological periods. the time scale is shown in millions of years ago (MYA). (D) Evolution-guided prioritization of 7,267 candidate EAPSs identified in rice. Each point represents one candidate EAPS. The x axis shows the site-level phyloP score calculated from the candidate EAPS codon, and the y axis shows the gene-level phyloP score calculated across the coding sequence. Candidates ranking in the top 10% for site-level acceleration and the top 10% for gene-level conservation were retained as high-confidence EAPSs. Red and orange points indicate high-confidence EAPSs; orange points denote high-confidence EAPSs in selected well-characterized defense-related genes. Blue points indicate candidates ranking in the top 10% for either site-level acceleration or gene-level conservation.

We next screened these multiple sequence alignments for sites where an inferred ancestral acidic residue was replaced by a phosphorylatable residue in Poaceae proteins. Ancestral acidic residues were inferred from conserved aspartate or glutamate residues in outgroups and early-diverging angiosperm lineages. Candidate EAPSs were defined as orthologous positions at which the computationally inferred ancestral acidic residues were replaced by serine, threonine or tyrosine in at least one of the five surveyed Poaceae species (Figure 1A). This analysis showed that such substitutions are common in grass proteomes. More than 15% of OGs in the sampled Poaceae species contain at least one candidate EAPS (Figure 1A). Across the five Poaceae species, we identified approximately 6,800 to 8,900 candidate EAPSs per species, spanning 4,800 to 6,200 genes (Figure 1B). In *Oryza sativa*, 7,267 candidate EAPSs were detected in 5,109 genes (Figure 1B). These results indicate that potential evolutionarily derived phosphoregulatory states are not sporadic, lineage-restricted events, but represent a broadly prevalent feature during grass protein evolution.

### Evolution-guided prioritization identifies high-confidence EAPSs in conserved rice genes

To prioritize high-confidence EAPSs for subsequent functional validation, we next used a substantially expanded, dense sampling of publicly available grass genomes to filter for sites that exhibit significant local sequence divergence within otherwise deeply conserved genes. We performed whole-genome alignments across 45 representative Poaceae species using the *O. sativa* genome as the reference (Figure 1C, Figure S3 and Table S2). For each of the 7,267 candidate EAPSs in rice, we used phyloP-based evolutionary metrics to quantify site-level evolutionary acceleration at each candidate codon and global gene-level conservation across the corresponding coding sequence (Figure 1D). Candidates ranked in the top 10% for both site-level acceleration and gene conservation were retained, yielding 151 high-confidence EAPSs distributed across 141 rice genes (Figure 1D and Table S3).

Among the 141 genes containing high-confidence EAPSs, nine are associated with rice immune-related processes, including *OsCPK12*, *OsMPK4*, *OsMPK3*, *OsATG7* and *OsRbohB*^16–20^ (Figure 1D). We selected *OsRbohB* for functional analysis because *OsRbohB* encodes a respiratory burst oxidase homologue that generates reactive oxygen species (ROS), serving as a central component of the plant immune response^20^.

### Immune-related rice Rboh proteins acquired phosphorylatable residues at a conserved EF-hand Ca^2+^-binding position

Among high-confidence candidate EAPSs, the Thr288 residue of OsRbohB resides within the second EF-hand motif in its cytosolic N-terminal regulatory region (Figure 2A). This residue maps precisely to the critical Y coordinate in the canonical EF-hand Ca^2+^ binding loop, a highly conserved position that is typically occupied by negatively charged acidic residues responsible for Ca^2+^ coordination^21^. Multiple sequence alignment of all nine rice Rboh proteins revealed that, among rice Rboh homologues, only the two immune-related paralogs OsRbohB and OsRbohI carry putative phosphorylatable residues at this functional site^20,22^, specifically Thr288 in OsRbohB and Ser317 in OsRbohI (Figure 2A and 2B).

**Figure 2.**
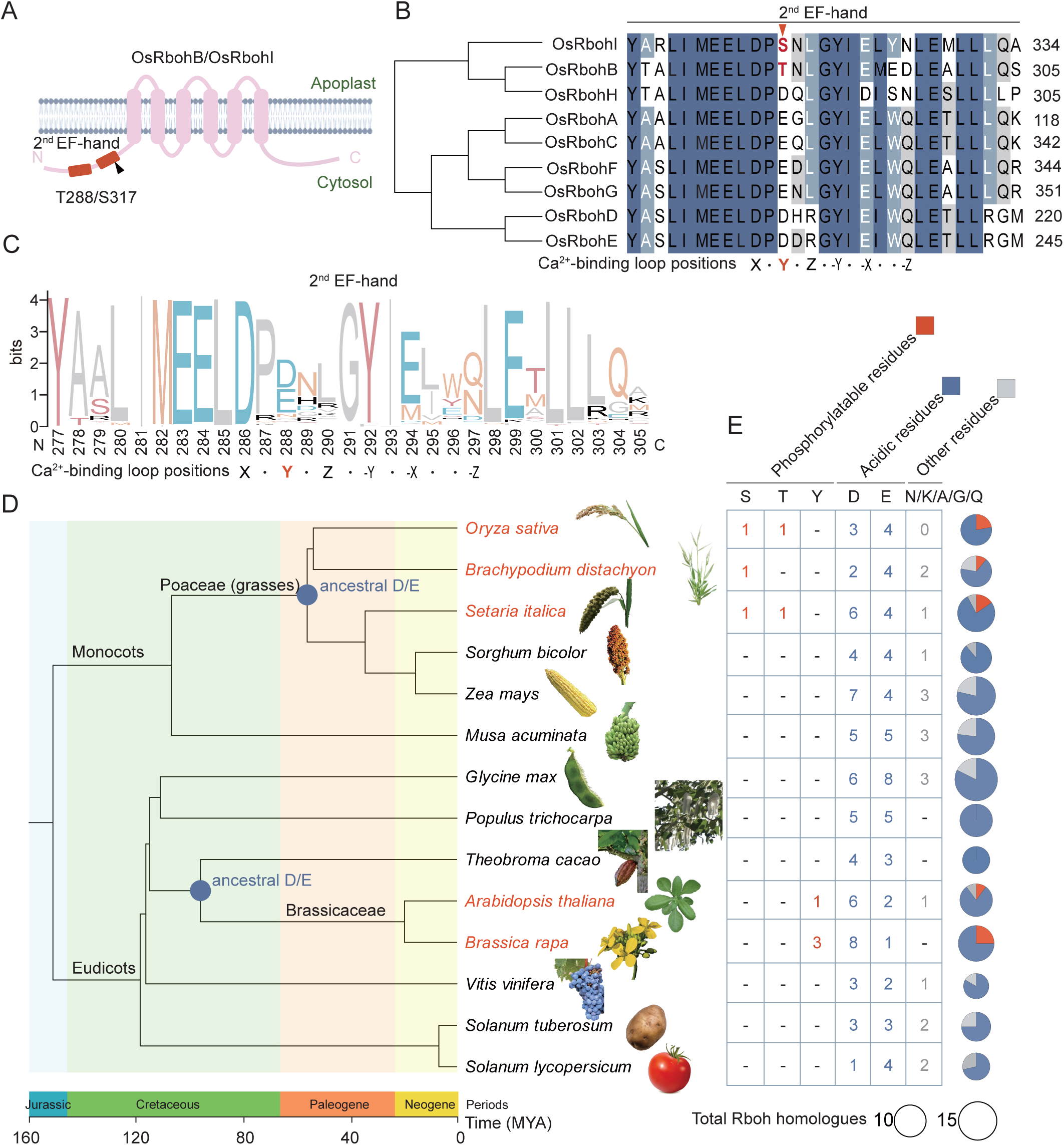
Immune-related rice Rbohs acquire phosphorylatable residues at a conserved EF-hand calcium-binding position. (A) Schematic representation of OsRbohB and OsRbohI topology, showing the N-terminal and cytosolic second EF-hand motif (in red) containing OsRbohB Thr288 and OsRbohI Ser317. (B) Phylogenetic relationship and sequence alignment of the second EF-hand motifs among the nine rice Rboh proteins. The Ca^2+^-binding loop is annotated by the canonical ligand positions X, Y, Z, −Y, −X and −Z. Phosphorylatable residues at the Y position, corresponding to Thr288 in OsRbohB and Ser317 in OsRbohI, are highlighted in red. (C) Sequence logo showing amino-acid frequencies within the second EF-hand Ca^2+^-binding loop across 167 Rboh homologues from 17 plant species. The Thr288-equivalent Y position is highlighted. (D and E) Phylogenetic mapping (D) and quantification (E) of residue states at the Y position of the second EF-hand Ca^2+^-binding loop across representative monocot and eudicot Rboh proteins. Blue circles in (D) indicate inferred ancestral acidic residues Asp or Glu. Species names in red indicate that the corresponding lineages harbor phosphorylatable residues at this position. The bottom color bar denotes geological periods, and the timescale is shown in million years ago (MYA). In (E), numbers indicate the counts of Rboh homologues carrying Ser, Thr, Tyr, Asp, Glu or other residues at this position. Pie charts summarize residue classes: phosphorylatable residues Ser, Thr or Tyr; acidic residues Asp or Glu; and other residues. Circle size indicates the total number of Rboh homologues analysed for each species.

Subsequently, we performed ancestral state reconstruction on a dataset comprising 167 Rboh homologues from 17 representative plant species to infer the evolutionary origin of these putative phosphorylatable residues. Strikingly, the position equivalent to OsRbohB Thr288 is predominantly occupied by acidic residues aspartate or glutamate across plant Rbohs (Figure 2C). Ancestral state reconstruction supports an ancestral acidic state at this critical EF-hand position, with derived phosphorylatable residues appearing in later-diverging lineages (Figure 2D and 2E). Specifically, serine or threonine residues were acquired in Poaceae Rbohs, including OsRbohB and OsRbohI, whereas tyrosine substitutions arose independently in Brassicaceae Rbohs. Thus, the conserved EF-hand Ca^2+^-binding motif has undergone independent functional remodeling across divergent plant lineages, transitioning from the ancestral acidic residue to derived phosphorylatable residues. These findings identify OsRbohB Thr288 and OsRbohI Ser317 as evolutionarily derived candidate phosphoregulatory sites in immunity-related rice NADPH oxidases.

### OsCPK17 phosphorylates OsRbohB at Thr288 and ancestral residue restoration enhances OsRbohB activity

We next investigated whether the OsRbohB Thr288 residue is phosphorylated during immune signaling and whether reverting this derived residue to its ancestral acidic state would alter OsRbohB activity. Given that *Arabidopsis* subgroup-I CPK5 phosphorylates RbohD to promote ROS production^23^, we tested whether OsCPK17, a rice subgroup-I CPK implicated in immune homeostasis^24^, functions as a direct kinase for OsRbohB. GST pull-down assays demonstrated that MBP-OsCPK17 was specifically retained by GST-OsRbohB^1–355^, but not by the GST control. Furthermore, co-immunoprecipitation assays in transfected rice protoplasts confirmed the *in vivo* association between OsCPK17-FLAG and HA-OsRbohB (Figure S4A and S4B).

*In vitro* phosphorylation assays showed that purified MBP-OsCPK17 directly phosphorylated the cytosolic N-terminal fragment of OsRbohB (His-OsRbohB^1-3^^55^). Notably, this phosphorylation signal was substantially attenuated by the T288A substitution, identifying Thr288 as a major OsCPK17-targeted phosphosite in this N-terminal region (Figure 3A and 3B). In rice protoplasts, OsRbohB phosphorylation was readily detected in wild-type cells, but was markedly reduced in *oscpk17*, supporting an OsCPK17 requirement for efficient phosphorylation of OsRbohB *in vivo* (Figure 3C). Using a custom-generated phospho-specific antibody against the Thr288-phosphorylated epitope of OsRbohB, we further demonstrated that both flg22 and chitin induced Thr288 phosphorylation in wild-type rice cells, whereas this induced phosphorylation was greatly reduced in *oscpk17* mutants and completely abolished in OsRbohB^T288A^ variant (Figure 3D). Collectively, these results confirmed that Thr288 is a phosphosite directly targeted by OsCPK17.

**Figure 3.**
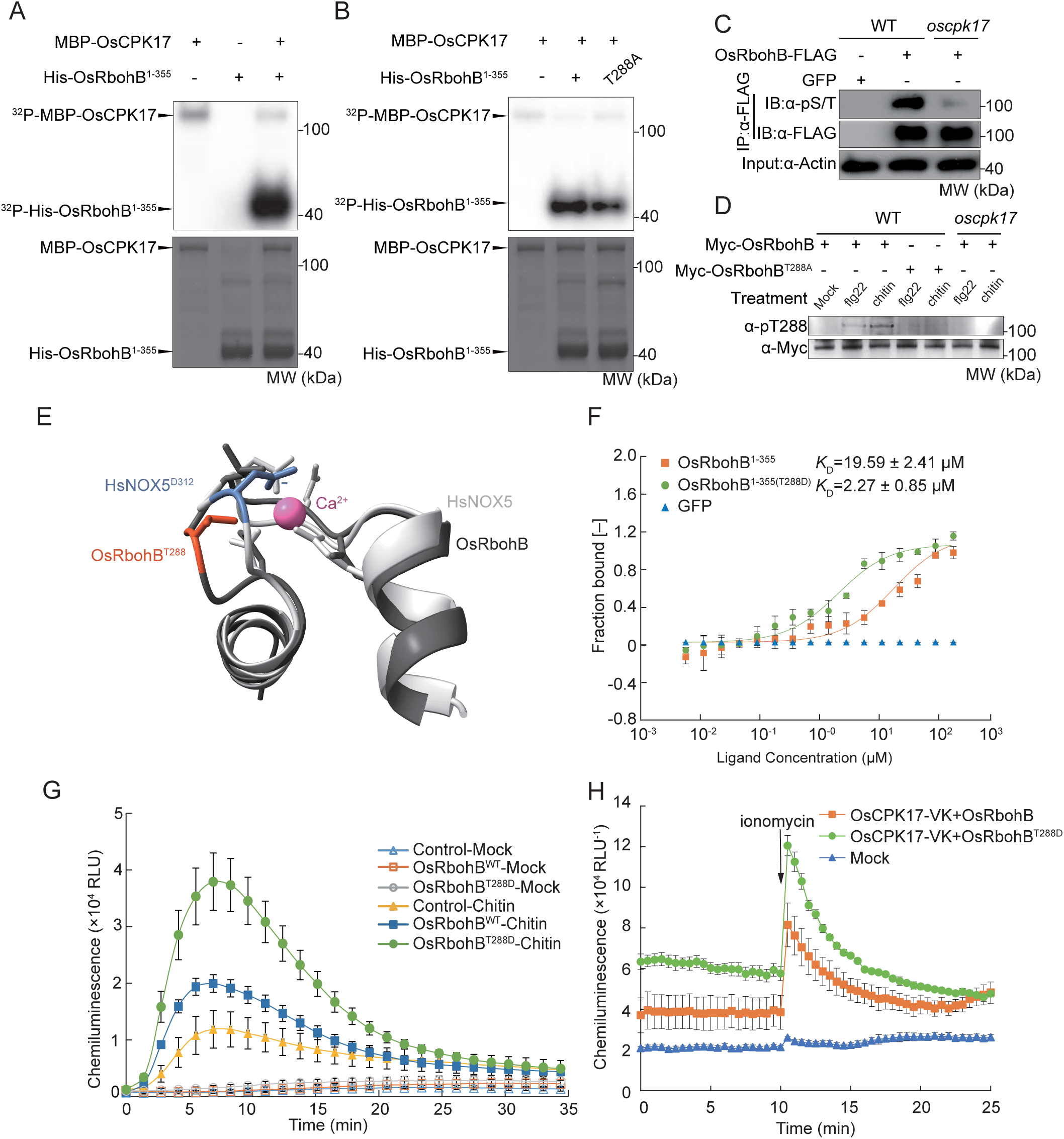
Restoration of the ancestral acidic residue at OsRbohB Thr288 enhances Ca^2+^ binding and ROS production. (A and B) *In vitro* kinase assays showing phosphorylation of the OsRbohB N-terminal cytosolic region by OsCPK17. Recombinant MBP-OsCPK17 was incubated with His-OsRbohB^1–3^^55^ or its T288A variant as indicated. Phosphorylation was detected by autoradiography, and protein loading was assessed by Coomassie brilliant blue staining. (C) *In vivo* phosphorylation of OsRbohB in rice protoplasts. OsRbohB-FLAG was transiently expressed in wild-type or *oscpk17* protoplasts and immunoprecipitated with anti-FLAG beads. Phosphorylation was detected using an anti-phosphoserine/threonine antibody, and OsRbohB-FLAG abundance was detected with an anti-FLAG antibody. Actin was detected by immunoblotting with anti-Actin antibody as the input loading control. (D) PAMP-induced phosphorylation of OsRbohB at Thr288 was detected using a phospho-specific anti-pT288 antibody. Myc-OsRbohB or Myc-OsRbohB^T288A^ was transiently expressed in wild-type or *oscpk17* rice protoplasts and treated with mock, chitin or flg22. Myc-OsRbohB abundance was detected with an anti-Myc antibody. (E) Structural comparison of the modelled OsRbohB EF-hand region with the human NOX5 Ca^2+^-binding structure. OsRbohB Thr288 and the corresponding human NOX5 Asp312 are involved in Ca^2+^ coordination. (F) Microscale thermophoresis analysis to show Ca^2+^ binding affinities of purified GFP, OsRbohB^1–3^^55^ and OsRbohB^1–3^^55(T288D)^ proteins. Curves show one-site binding fits; apparent dissociation constants (*K*_D_) are shown as means ± s.e.m.; n = 3. (G) Chitin-induced ROS burst in *N. benthamiana* leaves transiently expressing GFP, OsRbohB^WT^-GFP or OsRbohB^T288D^-GFP. ROS production was measured by luminol-based chemiluminescence. Data are shown as means ± s.e.m.; n = 8. (H) Ionomycin-induced ROS production in HEK293T cells expressing OsRbohB^WT^ or OsRbohB^T288D^ together with OsCPK17-VK, a 1–400-aa fragment lacking the C-terminal EF-hand-containing regulatory domain. Ionomycin was added at the indicated time point. Data are mean ± s.e.m.; n = 4.

Structural modelling of OsRbohB, compared with the human NOX5-Ca^2+^ complex structure, locates Thr288 in the calcium-binding loop of the second EF-hand, with a spatial arrangement that supports its involvement in Ca^2+^ coordination (Figure 3E). Because Ca^2+^ binding to N-terminal EF-hand motifs has been well documented as an important activation mechanism for plant Rbohs^25^, we tested whether restoration of the ancestral acidic residue at this position alters Ca^2+^ binding affinity and oxidase activity. Microscale thermophoresis assays showed that OsRbohB^1-3^^55(T288D)^ exhibited higher Ca^2+^ binding affinity than wild-type OsRbohB^1-3^^55^, with apparent dissociation constants of 2.27 ± 0.85 μM and 19.59 ± 2.41 μM, respectively (Figure 3F). A parallel ancestral residue substitution S317D in OsRbohI^1-383^ also increased Ca^2+^ binding affinity, reducing the apparent dissociation constant from 16.8 ± 2.4 μM to 2.3 ± 0.8 μM (Figure S5A). Consistent with the enhanced Ca^2+^ binding affinity, OsRbohB^T288D^ and OsRbohI^S317D^, when transiently expressed in *Nicotiana benthamiana*, triggered significantly stronger chitin-induced reactive oxygen species (ROS) bursts than their respective wild-type proteins (Figure 3G and Figure S5B). Similarly, we employed HEK293T cells, a standard heterologous reconstitution system lacking endogenous plant CPK/Rboh homologs, to test activation of OsRbohB. In cells co-expressing OsRbohB variants together with OsCPK17-VK, a truncated OsCPK17 protein comprising residues 1–400 and lacking the C-terminal EF-hand-containing regulatory domain, OsRbohB^T288D^ generated significantly more ROS than wild-type OsRbohB following ionomycin treatment (Figure 3H). Collectively, these findings indicate that restoring ancestral acidic residues in the EF-hand motifs enhances Ca^2+^-dependent activation of immune-related rice Rbohs, and further identify OsRbohB Thr288 as a key residue linked to OsCPK17-dependent immune phosphorylation.

### Restoration of the ancestral state of OsRbohB via prime editing enhances rice immunity

We next tested whether introducing the T288D substitution via targeted editing of OsRbohB enhances rice disease resistance. Through prime editing, we generated two independent *OsRbohB^T288D^* lines in the Zhonghua 11 (ZH11) background, in which the endogenous Thr288 codon was precisely edited to encode Asp (Figure S6). Upon treatment with chitin and flg22, both edited lines produced more robust ROS bursts than wild-type plants (Figure 4A and 4B). In line with this result, expression of the defense marker genes *OsPBZ1* and *OsPAL1* was significantly higher in *OsRbohB^T288D^*plants after chitin or flg22 elicitation (Figure 4C, 4D, S7A and S7B). We therefore conclude that restoring the ancestral residue at OsRbohB Thr288 potentiates pathogen-associated molecular pattern (PAMP)-triggered immunity in rice.

**Figure 4.**
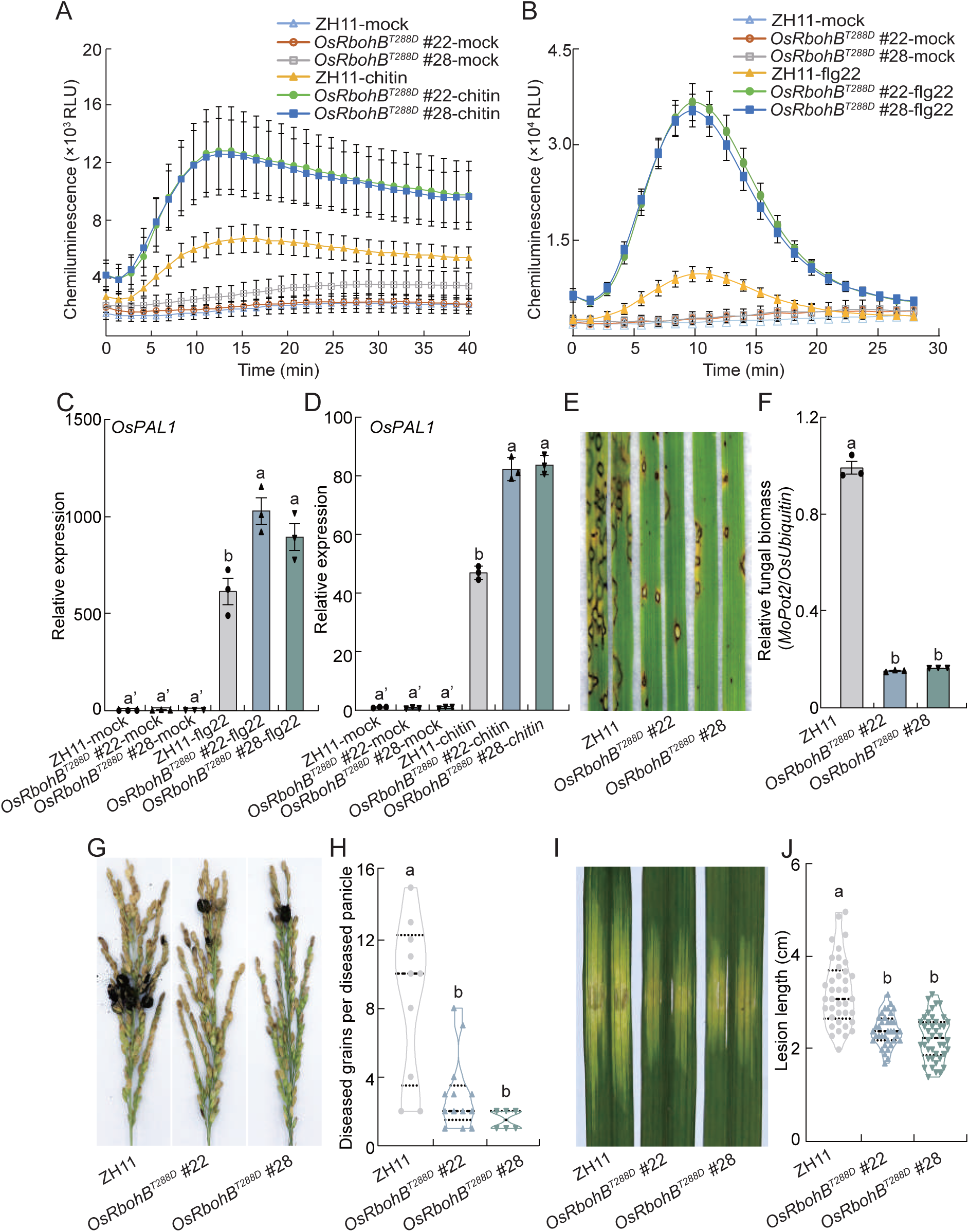
Restoration of the ancestral acidic residue at OsRbohB Thr288 enhances PAMP-triggered immunity and broad-spectrum disease resistance in rice. (A and B) Chitin-induced (A) and flg22-induced (B) ROS bursts in leaves of wild-type ZH11 and two independent *OsRbohB^T288D^* prime-edited lines. Data are mean ± s.e.m.; n = 16. (C and D) Relative expression of *OsPAL1* in ZH11 and *OsRbohB^T288D^* plants after flg22 (C) or chitin (D) treatment. Transcript levels were normalized to *OsActin1*. Data are mean ± s.e.m.; n = 3. Different letters indicate statistically significant differences in gene expression level. (E and F) Representative rice blast lesions (E) and relative fungal biomass (F) in ZH11 and *OsRbohB^T288D^* plants after inoculation with *M. oryzae* isolate RB22. Fungal biomass was quantitatively determined using a DNA-based qPCR assay, where the *M. oryzae*-specific transposable element *MoPot2* and the rice *Ubiquitin* reference gene were employed as the pathogen and host plant genomic markers, respectively. Data are mean ± s.e.m.; n = 3. (G and H) Representative rice false smut symptoms (G) and quantification of diseased grains per diseased panicle (H) after inoculation with *U. virens* strain U14Kc002. Horizontal lines indicate the median and quartiles. Different letters indicate statistically significant differences. (I and J) Representative bacterial leaf streak symptoms (I) and lesion-length quantification (J) after inoculation with *X. oryzae* pv. *oryzicola* strain

We then examined whether this heightened immune activation translates to enhanced disease resistance. Upon *Magnaporthe oryzae* inoculation, *OsRbohB^T288D^*lines exhibited reduced blast lesion sizes and substantially less fungal biomass compared with wild-type plants (Figure 4E and 4F). The edited lines also showed enhanced resistance to rice false smut caused by *Ustilaginoidea virens*, with fewer diseased grains on inoculated panicles than wild-type plants (Figure 4G and 4H). In addition, *OsRbohB^T288D^*plants displayed shorter disease lesions after inoculation with *Xanthomonas oryzae* pv. *oryzicola*, indicating increased resistance to bacterial leaf streak (Figure 4I and 4J). Collectively, these results demonstrate that restoring the ancestral acidic residue at OsRbohB Thr288 increases broad resistance to fungal and bacterial diseases in rice.

We further investigated whether this functional pattern applies to OsRbohI Ser317. We employed prime editing to generate *OsRbohI^S317D^*mutant lines, where the native Ser317 codon was precisely edited to encode aspartate (Figure S8). Similar to *OsRbohB^T288D^* plants, *OsRbohI^S317D^* edited lines showed enhanced chitin- and flg22-induced ROS bursts (Figure S9A and S9B). Chitin- or flg22-triggered expression of *OsPBZ1* and *OsPAL1* was also higher in *OsRbohI^S317D^* plants than in wild-type plants (Figure S9C–F). These lines exhibited increased resistance to *M. oryzae* and *X. oryzae* pv. *oryzicola* infection (Figure S9G-J). These results indicate that ancestral-state restoration at equivalent EF-hand positions in two immune-related rice Rbohs boosts rice immunity and confers enhanced resistance against multiple pathogens.

We further evaluated whether the enhanced OsRbohB-mediated immunity incurs any yield-related trade-offs under field conditions. Phenotypic analyses of mature plants and harvested grains showed that the *OsRbohB^T288D^* lines displayed no detectable alterations in overall plant architecture, thousand-grain weight, plant height, grain length, or grain width compared with the wild type (Figure S10A–F). Effective tiller number was even increased in the edited lines (Figure S10G). Taken together, these findings indicate that restoration of the ancestral acidic residue in OsRbohB enhances rice immunity and broad-spectrum disease resistance without compromising major yield-related traits under the fertilized conditions tested.

### Field testing infer loss of the ancestral phospholatable residues due to growth-defense trade-offs in nutrient-limiting condition

Restoring the ancestral acidic residue at OsRbohB Thr288 enhances immune output without obvious fitness cost, raising the question of why the ancestral acidic state was replaced by a phosphorylatable residue during rice Rboh evolution. We next traced the evolutionary origin of the Thr288 residue in OsRbohB by examining the corresponding orthologous site across wild and cultivated rice relatives. The phosphorylatable Thr is shared by cultivated *O. sativa* and multiple wild *Oryza* species, including *O. rufipogon*, *O. nivara* and *O. barthii* (Figure 5A). Residue mapping indicates that the Asp-to-Thr substitution event occurred before the diversification of these wild and cultivated rice lineages. Additionally, a broad survey of geographically and genetically diverse cultivated rice accessions showed that Thr288 is fixed across the cultivated subpopulations examined, including both japonica and indica rice (Figure 5B). Thus, phylogenetic reconstruction indicates that the Thr288 residue predated rice domestication, rather than evolving as a newly derived allele during rice domestication.

**Figure 5.**
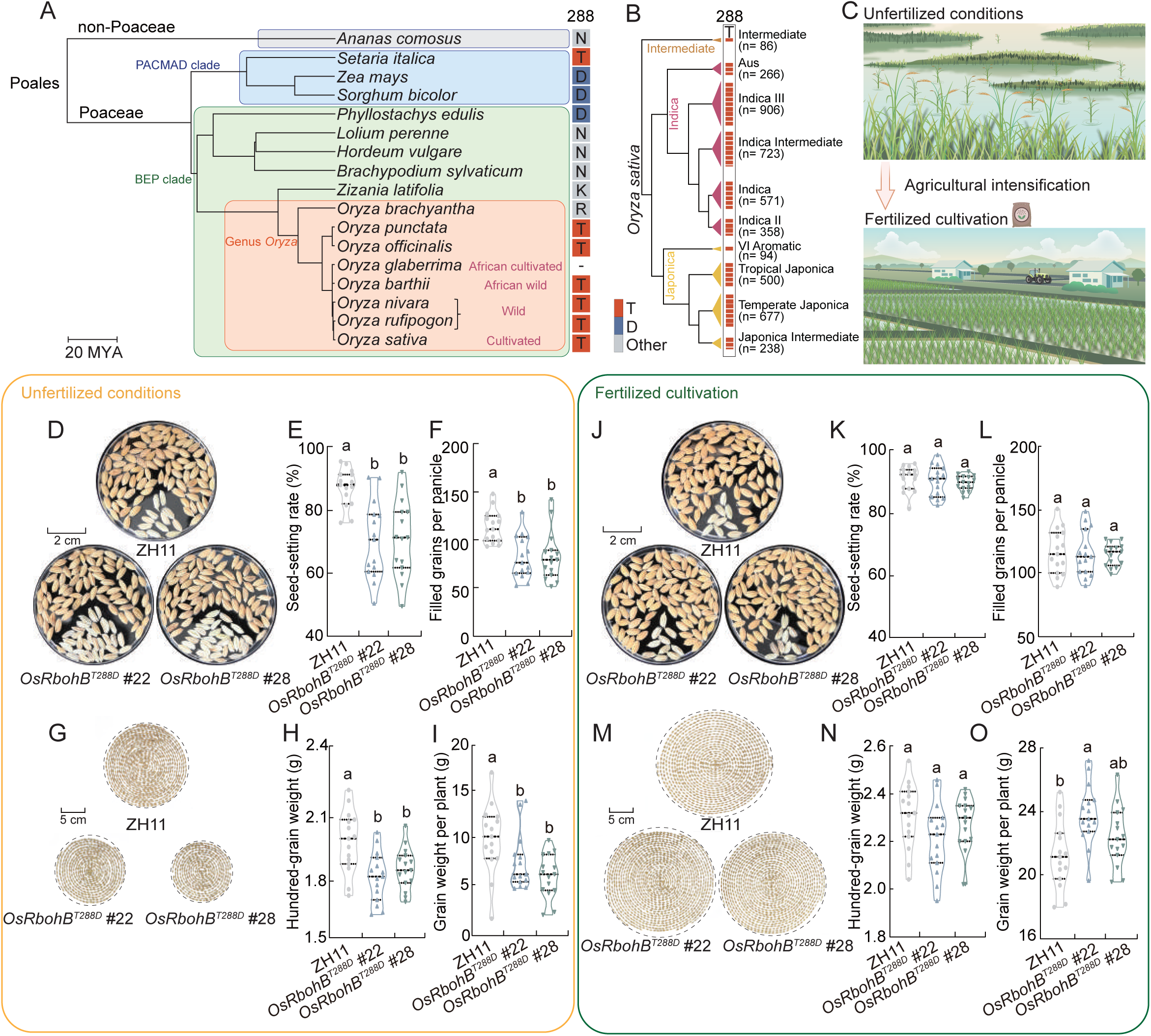
The growth–defense trade-off caused by ancestral-state restoration at OsRbohB Thr288 is alleviated under fertilized cultivation. (A) Phylogenetic distribution of amino-acid residues at the OsRbohB Thr288-equivalent position across representative Poales species. The tree includes a non-Poaceae outgroup, PACMAD and BEP clade species, the two major Poaceae clades, and representative *Oryza species*. Letters on the right indicate the amino-acid residues at the Thr288-equivalent position. No OsRbohB orthologue was found in *Oryza glaberrima*. Scale bar, 20 million years. (B) Thr288 is invariant across the examined cultivated rice accessions. Amino-acid residues at OsRbohB Thr288 across 4,419 cultivated rice accessions were obtained from RiceVarMap2. The population tree follows the RiceVarMap2 subpopulation classification. Bars indicate accession numbers in each subpopulation, with sample sizes shown in parentheses. (C) Conceptual illustration of the contrast between unfertilized natural environments experienced during rice evolution and modern rice cultivation supported by irrigation and fertilizer inputs. (D-I) Yield-related agronomic traits of wild-type and two independent *OsRbohB^T288D^* prime-edited lines in the paddy field without fertilizer application. (D) Representative images of filled and unfilled grains to assess seed setting. (E) Seed-setting rate. (F) Filled grains per panicle. (G) Representative images of harvested grains per plant. (H) Hundred-grain weight. (I) Grain weight per plant. Scale bars, 2 cm (D) and 5 cm (G). (J-O) Yield-related agronomic traits of wild-type and *OsRbohB^T288D^* lines under fertilized cultivation. (J) Representative images of filled and unfilled grains to assess seed setting. (K) Seed-setting rate. (L) Filled grains per panicle. (M) Representative images of harvested grains per plant. (N) Hundred-grain weight. (O) Grain weight per plant. Scale bars, 2 cm (J) and 5 cm (M). For (E, F, K and L), n = 15 plants. For (H, I, N and O), n = 15 plants. Violin plots show the distribution of individual values; horizontal lines indicate the median and quartiles. Different letters indicate statistically significant differences among different rice lines.

Wild rice populations evolved across heterogeneous natural habitats, where nutrient availability is likely limited in some habitats, whereas modern rice cultivation largely relies on irrigation and fertilizer inputs (Figure 5C). Two competing models for growth-defense tradeoffs in plants are based on “external resource ceiling” vs. “internal genetic wiring”^26^. We hypothesized that the OsRbohB ancestral Asp288 allele may incur a fitness cost under resource-limited ancestral habitats, which enables a rigorous field-based evaluation of the two competing models underlying growth-defense tradeoffs. To test this hypothesis, we conducted a side-by-side comparative assay in a previously uncultivated field in Hainan, China, to evaluate the fitness cost associated with restoring the ancestral acidic residue under nutrient-poor (mimicking hypothetical ancestral habitats) and nutrient-rich conditions (representing modern rice cultivation systems) (Figure S11). Indeed, under unfertilized cultivation conditions, the *OsRbohB^T288D^* edited lines showed reduced reproductive performance compared with wild-type plants, including decreases in seed-setting rate, filled grains per panicle, hundred-grain weight and grain weight per plant (Figure 5D–I), with no marked reduction in vegetative growth traits (Figure S12A and S12B). Thus, restoring the ancestral acidic state imposes a measurable reproductive fitness cost under nutrient-limited conditions, indicating that the derived Thr288 variant mitigates this fitness penalty in low-nutrient environments.

Under standard fertilized cultivation, by contrast, *OsRbohB^T288D^* plants showed no visible reduction in seed production relative to wild-type plants (Figure 5J). Quantification of yield-related traits detected no significant differences in seed-setting rate, filled grains per panicle or hundred-grain weight between the wild-type and edited lines (Figure 5K–N). Grain weight per plant showed no reduction and was even increased in one edited line (Figure 5O), while vegetative growth traits remained unaltered (Figure S12C and S12D). Collectively, the results indicate that reproductive cost of restoring the ancestral OsRbohB state is effectively buffered under modern fertilized cultivation regimes.

## Discussion

Our study demonstrates a potentially paradigm-shifting concepts with significant broad impacts. We show that evolutionary history can be harnessed as a recoverable reservoir of untapped molecular variation for crop improvement. In sharp contrast, conventional breeding has long relied almost exclusively on alleles preserved in cultivated germplasm, landraces and wild relatives as the core genetic materials^1,2^. This strategy has been highly productive, but it is constrained by the limited fraction of standing historical variation that remains accessible in living populations. Acidic-to-phosphorylatable residue substitutions have been recognized as an evolutionary route for the de novo emergence of functional phosphorylation sites, yet this type of mutations has rarely been explored as a source of functional variation for modern breeding^10^. Here, by identifying candidate EAPSs across grasses and restoring ancestral acidic residues in rice Rboh proteins, we demonstrate that historical residue states lost during evolution and absent from contemporary crop variation can be inferred from protein evolution, rebuilt by precision editing, and converted into elite alleles that enhance immune output and broad-spectrum disease resistance (Figure 1-4). To our knowledge, this represents the first experimental demonstration that historical molecular variation, rather than only standing genetic variation, can be directly used for modern crop improvement. This ancestral-state restoration strategy therefore extends crop improvement beyond mining existing cultivated alleles, establishing a new paradigm for exploiting deep evolutionary history as an untapped breeding resource.

EAPSs represent a largely uncharacterized class of candidate functional sites that directly bridge long-term protein evolution to rewired regulatory functions. As one of the most pervasive mechanisms of post-translational regulation, protein phosphorylation acts as a central regulatory layer governing plant immune signalling^27–30^. Acidic-to-phosphorylatable substitutions are uniquely tractable: they create a single-residue contrast between a constitutively charged ancestral state and a derived phosphorylation-dependent state^10^. Here, we integrated sequence-evolution analysis with ancestral-state reconstruction, prioritizing candidate codons with local evolutionary acceleration that reside within broadly conserved coding sequences (Figure 1). This contrast between site-level regulatory remodeling and gene-level functional constraint precisely pinpoints residues where historical regulatory states can be computationally reconstructed. In crop plants, candidate EAPSs were identified in all five sampled Poaceae species, laying a solid basis for evaluating this approach beyond rice (Figure 1A and 1B). Functional tests of six candidate EAPSs across five rice genes implicated in plant immunity further confirmed that restoring the inferred ancestral residue at four of these sites enhanced immune responses (data not shown). Together, these findings suggest that ancestral-state restoration may be applicable to additional immune-related genes and other grass species. For crop improvement, such single-residue precision edits can generate targeted regulatory elite alleles, while avoiding the linkage drag associated with introgressing large segments from distant exotic germplasm.

Our study also provides strong field-based evidence that the classic growth-defense trade-off, which is likely shaped under ancestral nutrient-limited conditions, can be fully uncoupled in modern nutrient-rich agricultural systems. Growth-defense trade-offs represent a long-standing bottleneck for breeding disease-resistant crop germplasm, because enhanced immunity often incurs penalties on plant growth or reproduction^11,12,26^. In OsRbohB, restoring the ancestral acidic residue at Thr288 enhanced immune ROS production and broad-spectrum disease resistance (Figure 3 and 4), but incurred a significant reproductive cost under nutrient-limited field conditions (Figure 5D–I). Under fertilized cultivation, this fitness cost was buffered, enabling the ancestral-state allele to enhance immunity without yield penalty (Figure 5J–O). Thus, the derived phosphorylatable Thr288 state was probably retained during evolution because it effectively mitigates the fitness cost of immune activation in ancestral nutrient-limited environments, whereas modern high-fertility cultivation systems relax this ancient evolutionary constraint, unlocking the hidden agronomic value of the ancestral acidic residue state.

We propose the term “Beneficial Ancestral Genomic Sites (BAGS)” to describe a class of historical molecular residues that were once constrained, replaced or disfavored under past ancestral environments, but gain high functional agronomic value when precisely reconstructed under modern agricultural contexts. The term encapsulates the core principle that a residue state once eliminated by one selective regime can become beneficial under a new environmental setting. In crops, this principle can extend beyond disease resistance. Modern agriculture has drastically reshaped nutrient availability, planting density, water management, genetic uniformity and biotic stress exposure, completely rewiring the selective context for optimizing critical traits ranging from immunity, stress tolerance, growth architecture, reproductive output to resource-use efficiency^1,11,13,31,32^. More broadly, the BAGS framework is applicable to any scenario where organisms experience rapid shifts in ecological, genetic or human-engineered environments, including domestication, artificial selection, host jumps, chemical pressure or climate stress. This paradigm establishes a conceptual framework to systematically explore how evolutionary history harbors hidden, recoverable molecular states that are invisible within modern standing variation, yet can unlock renewed functional value under contemporary environmental conditions.

## STAR METHODS

### Key resources table

| REAGENT or RESOURCE | SOURCE | IDENTIFIER |
|---|---|---|
| **Antibodies** |  |  |
| Anti-FLAG M2 antibody | Sigma-Aldrich | F1804 |
| HRP-conjugated anti-HA antibody | Roche | 12013819001 |
| Anti-phosphoserine/threonine antibody | Abcam | AB117253 |
| Anti-plant actin antibody | LABLEAD | BP0101 |
| OsRbohB phospho-Thr288-specific antibody | Abmart | N/A |
| Anti-Myc antibody | ABclonal | AE010 |
| Anti-His antibody | CWBIO | CW0286M |
| Anti-GST antibody | LABLEAD | G1001 |
| **Chemicals, peptides, and recombinant proteins** |  |  |
| Anti-FLAG M2 affinity gel | Sigma-Aldrich | A2220 |
| HisSep Ni-NTA Agarose Resin | YEASEN | 20502ES60 |
| GSTSep Glutathione Agarose Resin 4FF | YEASEN | 20508ES60 |
| Protease inhibitor cocktail (100×) | CWBIO | CW2200S |
| Cellulase “Onozuka” R-10 | Yakult Pharmaceutical Industry Co., Ltd. | 260508-02 |
| Macerozyme R-10 | Yakult Pharmaceutical Industry Co., Ltd. | 250710-02 |
| L-012 | FUJIFILM Wako Pure Chemical Corporation | 120-04891 |
| Horseradish peroxidase | Coolaber | CH6161 |
| Chitin from shrimp shells | Sigma-Aldrich | C9752 |
| flg22 peptide | Sangon Biotech | N/A |
| Ionomycin calcium salt, ready-made solution | Sigma-Aldrich | I3909 |
| Polyethylenimine (PEI) | XANCON | CK012 |
| Potassium-sulfate-type compound fertilizer (15–15–15, N–P<sub>2</sub>O<sub>5</sub>–K<sub>2</sub>O) | Kingenta Ecological Engineering Group Co., Ltd. | N/A |
| **Critical commercial assays** |  |  |
| BCA Protein Assay Kit | Coolaber | SK1070 |
| M5 PLANTpure plant total RNA extraction kit | Mei5bio | MF456 |
| M5 Super Plus qPCR RT Kit with gDNA Remover | Mei5bio | MF166-plus |
| 2× M5 HiPer Realtime PCR Super Mix with Low ROX (SYBR Green, with anti-Taq) | Mei5bio | MF797 |
| Ultra-sensitive ECL chemiluminescent substrate | Biosharp | BL520B |
| **Experimental models: cell lines** |  |  |
| HEK293T | Zhengxin Ying laboratory | RRID: CVCL_0063 |
| **Experimental models: organisms/strains** |  |  |
| *Oryza sativa* subsp. *japonica* cv. Dongjin (DJ) | laboratory collection | N/A |
| *Oryza sativa* subsp. *japonica* cv. Zhonghua 11 (ZH11) | laboratory collection | N/A |
| *Oryza sativa oscpk17* (DJ background) | POSTECH Rice Insertion Mutant Library; Mou et al.<sup>24</sup> | 4A-50582 |
| *Oryza sativa* OsRbohB T288D edited lines (ZH11 background) | This paper | LOC_Os01g25820.1; OsRbohB T288D |
| *Oryza sativa* OsRbohI S317D edited lines (ZH11 background) | This paper | LOC_Os11g33120.1; OsRbohI S317D |
| *Nicotiana benthamiana* | laboratory collection | N/A |
| **Recombinant DNA** |  |  |
| pET28a | Laboratory collection | Table S5 |
| pUC19-35S-FLAG-RBS | Jianmin Zhou's laboratory | Table S5 |
| pRTV-nHA | He et al.<sup>45</sup> | Table S5 |
| pRTV-nMyc | He et al.<sup>45</sup> | Table S5 |
| pRTV-GFP | He et al.<sup>45</sup> | Table S5 |
| pGEX-4T-1 | Laboratory collection | Table S5 |
| pGEX-4T-3 | Laboratory collection | Table S5 |
| pGD-GFP | Tao Zhou's laboratory | Table S5 |
| pEF1-2xStrepII-C-term | Jörg Kudla's laboratory | Table S5 |
| pEF1-2xStrepII-N-term | Jörg Kudla's laboratory | Table S5 |
| Modified PE-P3 prime-editing vector | Xu et al.<sup>58</sup> | Method details |
| OsRboh and OsCPK17 expression constructs | This paper | Method details; Table S4 |
| **Software and algorithms** |  |  |
| BLAST+ | Camacho et al.<sup>59</sup> | https://blast.ncbi.nlm.nih.gov/ |
| markov_clustering | Guy Allard | https://github.com/GuyAllard/markov_clustering |
| MAFFT v7.487 | Katoh and Standley<sup>36</sup> | https://mafft.cbrc.jp/alignment/software/ |
| Mashtree | Katz et al.<sup>37</sup> | https://github.com/lskatz/mashtree |
| Mash | Mash developers | https://github.com/marbl/Mash |
| RepeatMasker | RepeatMasker developers | https://www.repeatmasker.org/ |
| Progressive Cactus | Armstrong et al.<sup>38</sup> | https://github.com/ComparativeGenomicsToolkit/cactus |
| phyloP (PHAST) | Hubisz et al.<sup>39</sup> | https://github.com/CshlSiepelLab/phast |
| MEGA 7.0 | Kumar et al.<sup>40</sup> | https://www.megasoftware.net/ |
| WebLogo | Crooks et al.<sup>42</sup> | https://github.com/WebLogo/weblogo |
| AlphaFold 3 | Abramson et al.<sup>48</sup> | https://github.com/google-deepmind/alphafold3 |
| UCSF ChimeraX | Pettersen et al.<sup>50</sup> | https://www.cgl.ucsf.edu/chimerax/ |
| Biostrings | Bioconductor | https://bioconductor.org/packages/Biostrings/ |
| R | R Foundation for Statistical Computing | https://www.r-project.org/ |
| GraphPad Prism 8.0 | GraphPad Software | https://www.graphpad.com/ |
| Adobe Photoshop | Adobe | https://www.adobe.com/products/photoshop.html |
| Custom analysis scripts | This paper | https://figshare.com/s/d2a2360142abc059913d |

### RESOURCE AVAILABILITY

#### Lead contact

Further information and requests for resources and reagents should be directed to the lead contact, Wenxian Sun.

#### Materials availability

Materials generated in this study are available from the lead contact upon reasonable request.

#### Data and code availability

All data supporting the findings of this study are available in the main text and Supplementary Information. Genome assemblies and protein sequences used for comparative and evolutionary analyses were obtained from publicly available databases; the corresponding sources are listed in Table S2 and Table S6. Rice sequence-variation and subpopulation information were obtained from RiceVarMap2. Species relationships and divergence-time information were obtained from TimeTree. The human NOX5 Ca^2+^-binding structure used for structural comparison is available from the Protein Data Bank under accession code 6SZ5. No new genome, transcriptome or proteome sequencing datasets were generated in this study. Additional data are available from the corresponding authors upon request.

## AUTHOR CONTRIBUTIONS

W.S., S.Y.H., J.W., H.W. and Y.D. conceived the study. J.W., Shuo Liu, B.M., X.Z., G.Z. and L.Z. performed biochemical and molecular experiments. J.W., Shuo Liu, H.L., J.Y., F.X. and S.Z. performed field experiments and agronomic trait measurements. Dongyu Shi, C.Y., M.Z., Q.W. and M.Q. generated and identified experimental plant materials. S.W. performed the OsRbohB structural analysis. Danbing Shan, X.L., Shanlin Liu and Y.D. performed evolutionary analyses. H.W., J.W. and T.L. performed computational biology analyses. J.W. and W.S. wrote the original draft. W.S., S.Y.H., J.W., H.W. and Y.D. revised and finalized the manuscript. All authors reviewed and approved the manuscript.

## ACKNOWLEDGMENTS

We thank Suomeng Dong (Nanjing Agricultural University), Yong-Hwan Lee (Seoul National University), Guo-Liang Wang (The Ohio State University), Zuhua He (CAS Center for Excellence in Molecular Plant Science) and Detlef Weigel (Max Planck Institute for Biology Tübingen) for their insightful discussions and technical support. We are grateful to Feng Qin and Shengxue Liu (China Agricultural University) for providing the *Rboh* cDNA, and to Jörg Kudla (University of Münster) for sharing the pEF1-2xStrepII-C-term and pEF1-2xStrepII-N-term plasmids. We also thank Yule Liu (Tsinghua University) for assistance with the FlexStation 3 reader. We thank Hengjiu Eternal Genediting (Beijing) Technology Co., Ltd., Wen Xu, Jinxiao Yang, Qi-Jun Chen and Yu Lu (China Agricultural University) for support with precision genome editing, Tao Lin (China Agricultural University) for providing access to computational server resources used for data analysis, Guanghui Yang and Zhengxin Ying (China Agricultural University) for their support in HEK293T cell culture and Jingyu Xu (Heilongjiang Bayi Agricultural University) for sharing information on the Rboh protein family in plants.

This research was supported by the National Key R&D Program of China (2024YFD1200600 to J.W.), the National Natural Science Foundation of China (32673371 to J.W. and 32293241 to W.S.), the China Agricultural Research System (CARS-01 to W.S.), the Joint Research Program of State Key Laboratory of Agricultural and Forestry Biosecurity (No. SKLJRP2510 to J.W.), and by the 2115 Talent Development Program and Chinese Universities Scientific Fund of China Agricultural University to J.W.

## EXPERIMENTAL MODEL AND STUDY PARTICIPANT DETAILS

### Plant materials and growth conditions

The *oscpk17* T DNA insertion line in the *Oryza sativa* subsp. *japonica* cv. Dongjin (DJ) background was obtained from the POSTECH Rice Insertion Mutant Library (Republic of Korea)^24^. The genome-edited lines were generated in the *O. sativa* subsp. *japonica* cv. Zhonghua 11 (ZH11) background. Three-week-old and six-week-old rice seedlings grown in controlled growth chambers were used for *Magnaporthe oryzae* and *Xanthomonas oryzae* pv. *oryzicola* inoculation, respectively. Field-grown rice plants at the late-booting stage were used for *Ustilaginoidea virens* inoculation. *Nicotiana benthamiana* plants were grown in environmentally controlled growth chambers under a 16 h light/8 h dark photoperiod at 25 °C.

## METHOD DETAILS

All experimental assays in this study were repeated independently at least three times unless otherwise stated. All gene constructs were validated by sequencing. The primers used in this study are listed in Table S4 and the strains and plasmids are listed in Table S5.

### Identification and evolutionary analysis of phosphorylation sites

Genome assemblies and protein sequences from 17 representative plant species were obtained from publicly available databases (Table S6). These species included one gymnosperm (*Pinus taeda*), two early-diverging angiosperms (*Amborella trichopoda* and *Nymphaea colorata*), and representative eudicots and monocots. Among them, five Poaceae species, *Oryza sativa*, *Brachypodium distachyon*, *Setaria italica*, *Sorghum bicolor* and *Zea mays*, were used to identify grass-lineage substitutions. Phylogenetic relationships among these species were obtained from the TimeTree database^33^.

Orthologous groups (OGs) were identified using the gene-family annotation pipeline as described previously^34,35^. Briefly, pairwise protein similarities among all annotated proteins from the representative plant species were calculated using an all-by-all BLASTP search on a local server. The resultant similarity relationships were used to construct a graph, where nodes represent genes and edges connect gene pairs with detectable protein sequence similarity. Subsequently, gene clusters were inferred using the Markov clustering algorithm implemented in the Python *markov_clustering* package with default parameters except the inflation parameter, which was set to 2. Genes not assigned to any multi-gene cluster were treated as orphan genes and retained as single-member families. Gene families containing at least two proteins were retained as OGs for cross-species residue comparison, and protein sequences within each OG were aligned using MAFFT v7.487^36^. In total, 15,804 OGs met this criterion and were used for subsequent analyses.

Candidate evolutionarily acquired phosphorylation sites were identified from multiple protein sequence alignments by searching for positions at which an inferred ancestral acidic residue was replaced by a phosphorylatable residue (serine, threonine or tyrosine) in the Poaceae lineage. Aspartate and glutamate residues were considered ancestral acidic states when they were conserved in the gymnosperm and early-diverging angiosperm outgroups and were also supported by conservation in additional non-Poaceae lineages.

For each Poaceae species, candidate EAPSs were summarized at both the residue and gene-family levels. Residue-level counts represented the number of acidic-to-phosphorylatable residue substitutions detected in orthologous proteins. Gene-family-level counts represented the number of OGs containing at least one candidate EAPS in the corresponding species. The proportion of EAPS-containing OGs was calculated as the number of OGs containing at least one candidate EAPS divided by the total number of surveyed OGs in that species.

### Whole-genome alignments and phyloP-based prioritization of candidate EAPSs

A phylogenetic tree of 45 Poaceae species representing distinct genera (Table S2) was constructed using Mashtree based on Mash distances calculated from genome assemblies^37^. Repetitive sequences in each genome were soft-masked with RepeatMasker. The inferred phylogenetic tree and soft-masked assemblies were then used as input for Progressive Cactus to generate a whole-genome alignment^38^. All downstream analyses were performed using *Oryza sativa* as the reference genome.

Using the whole-genome alignment and inferred phylogeny, we calculated site-specific evolutionary conservation scores across the rice reference genomes with phyloP (mode = CONACC, method = GERP) ^39^. Gene-level conservation was defined as the mean phyloP scores calculated across all nucleotide positions within coding sequences (CDSs). For each putative phosphorylation site, evolutionary conservation was defined as the average phyloP score of the three nucleotides within the corresponding codon.

### Rice Rboh phylogenetic analysis and EF-hand alignment

Rice Rboh protein sequences were aligned using the ClustalW algorithm implemented in MEGA 7.0^40^. A phylogenetic tree was constructed using the Neighbor-Joining method with 1,000 bootstrap replicates to assess the reliability of the tree topology. The second EF-hand motifs of rice Rboh proteins were extracted from the multiple sequence alignment and used to compare the amino-acid residues at the Thr288-equivalent position.

### Cross-species Rboh sequence alignment and residue-state analysis

Sequences of 167 Rboh proteins from 17 plant species, spanning gymnosperms (*Pinus taeda*), basal angiosperms (*Amborella trichopoda*, *Nymphaea colorata*), monocots (*Musa acuminata*, *Brachypodium distachyon*, *Oryza sativa*, *Setaria italica*, *Zea mays*, *Sorghum bicolor*), and eudicots (*Vitis vinifera*, *Theobroma cacao*, *Brassica rapa*, *Arabidopsis thaliana*, *Populus trichocarpa*, *Glycine max*, *Solanum tuberosum*, *Solanum lycopersicum*), were retrieved from a previous study^41^. Protein sequences were aligned using the ClustalW algorithm implemented in MEGA 7.0^40^. The second EF-hand Ca^2+^-binding loop was extracted from the alignment, and amino-acid frequencies at each position were visualized using WebLogo^42^. Amino-acid residues at the Thr288-equivalent position were classified as phosphorylatable residues (Ser, Thr or Tyr), acidic residues (Asp or Glu), or other residues. The phylogenetic framework used for residue mapping was obtained from the TimeTree database^33^.

### Expression and purification of recombinant proteins

The CDS of *OsCPK17* was amplified using gene-specific primers and subcloned into a modified pET28a vector to express His_6_-MBP-OsCPK17, hereafter referred to as MBP-OsCPK17. The coding sequence corresponding to the N-terminal region of *OsRbohB* (*OsRbohB^1-3^*^55^) was cloned into pET28a and pGEX-4T-3 to express His-OsRbohB^1-3^^55^ and GST-OsRbohB^1-3^^55^, respectively.

The constructs were transformed into *Escherichia coli* BL21(DE3) (TransGen, Beijing, China) for protein expression. Cell cultures were grown to an OD_600_ of 0.6–0.8 and were then treated with 1 mM IPTG at 28 °C for 4 h to induce protein expression. His- and GST-tagged proteins were purified from bacterial lysates using Ni-NTA His·Bind Resin (Millipore, Burlington, MA, USA) and Glutathione-Sepharose beads (GE Healthcare, Chicago, IL, USA), respectively, according to the manufacturers’ instructions. Protein concentration was determined using a BCA Protein Assay Kit (Thermo Fisher Scientific, Waltham, MA, USA) with bovine serum albumin as the standard.

***In vitro* GST pull-down assay**

GST pull-down assays were performed as described with minor modifications^43^. Briefly, purified GST-OsRbohB^1-3^^55^ or GST alone (10 μg) was immobilized on 40 μl of GST·Bind Resin (Millipore, Burlington, MA, USA) and incubated with purified His_6_-MBP-OsCPK17 (10 μg) at 4 °C for 4 h in binding buffer (50 mM Tris-HCl, pH 7.4, 150 mM NaCl, 1 mM EDTA, and 1% Triton X-100). The resin was collected by centrifugation and washed thoroughly with ice-cold PBS. Bound proteins were eluted by boiling the resin in 100 μl SDS-PAGE loading buffer for 5 min. Eluted proteins were resolved by SDS-PAGE and detected by immunoblotting.

### Isolation and transformation of rice protoplasts

Twelve-day-old rice seedlings were cut into thin strips with a sterile surgical blade and transferred into a 150 ml sterile Erlenmeyer flask containing 20 ml enzyme solution consisting of 1.5% (w/v) Cellulase “Onozuka” R-10 and 0.75% (w/v) Macerozyme R-10 (Yakult Pharmaceutical Industry Co., Ltd., Tokyo, Japan), 0.6 M mannitol, 10 mM MES, pH 5.7, 1 mM CaCl_2_, 0.1% BSA and 0.035% β-mercaptoethanol (v/v). The samples were incubated at 28 °C in the dark with gentle shaking at 80 rpm for 4–6 h. Cell debris was removed by filtration through a 40-mesh sieve. The protoplasts were collected by centrifugation at 1,200 rpm for 5 min and washed twice with 40 ml W5 buffer (154 mM NaCl, 125 mM CaCl_2_, 5 mM KCl, and 2 mM MES, pH 5.7).

The protoplasts were resuspended in 1 ml of MMG buffer (0.4 M mannitol, 15 mM MgCl_2_, 4 mM MES, pH 5.7) and placed on ice for 30 min. For transfection, approximately 50 μg of plasmid DNA was mixed with 1 ml protoplast suspension (2×10^5^ cells ml^-1^) and 1.1 ml of PEG-CaCl_2_ solution (40% PEG 4000, 0.2 M mannitol and 100 mM CaCl_2_) in a 10 ml centrifuge tube, followed by incubation at 28 °C for 10 min. The transfection was terminated by adding 5 ml of W5 buffer. After centrifugation at 1,200 rpm for 5 min, the transfected protoplasts were resuspended in 5 ml of W5 buffer and incubated overnight at 25 °C in the dark.

### Co-immunoprecipitation assay

The CDSs of *OsRbohB* and *OsCPK17* were amplified using gene-specific primers and subcloned into pRTV-nHA and pUC19-35S-FLAG-RBS, respectively, to express HA-OsRbohB and OsCPK17-FLAG^44,45^. The constructs were transfected into rice protoplasts by PEG-mediated transformation. Total proteins were extracted from transfected protoplasts using IP buffer (50 mM Tris-HCl, pH 7.4, 150 mM NaCl, 1 mM EDTA, and 1% Triton X-100) supplemented with protease inhibitor cocktail (CWBIO, Beijing, China). Co-immunoprecipitation assays were performed as described previously with minor modifications^46^. Briefly, the protein extract (1 ml) was incubated with 25 µl anti-FLAG M2 affinity beads (Sigma-Aldrich, St. Louis, MO, USA) at 4 °C for 4 h with gentle rotation. The beads were washed thoroughly with ice-cold PBS. Bound proteins were eluted by heating the beads at 95 °C for 5 min in SDS-PAGE loading buffer. Input and immunoprecipitated proteins were resolved by SDS-PAGE and transferred onto polyvinylidene difluoride (PVDF) membranes. The membranes were probed with anti-FLAG antibody (Sigma-Aldrich, St. Louis, MO, USA; 1:5,000) and HRP-conjugated anti-HA antibody (Roche, Basel, Switzerland; 1:2,000). Signals were detected using an enhanced chemiluminescence detection system (Biosharp, Beijing, China).

### *In vitro* kinase assay

*In vitro* kinase assays were performed as previously described with minor modifications^47^. Briefly, test kinase protein (1 μg) and substrate protein (2 μg) were incubated in kinase assay buffer containing 25 mM Tris-HCl (pH 7.5), 10 mM MgCl_2_, 10 mM CaCl_2_, 1 mM DTT, 50 μM unlabelled ATP, and 1 μCi [γ-^32^P] ATP at 30 °C for 30 min. The reactions were terminated by adding SDS-PAGE loading buffer and heating at 95 °C for 5 min. Proteins were separated on 12.5% SDS-PAGE gels. After electrophoresis, protein phosphorylation was detected by autoradiography using a Storage Phosphor Screen and Typhoon Trio Variable Mode Imager (GE Healthcare, Chicago, IL, USA).

### *In vivo* phosphorylation assay

The coding sequence of *OsRbohB* was amplified using gene-specific primers and subcloned into pUC19-35S-FLAG-RBS. The construct was transfected into wild-type and *oscpk17* rice protoplasts to express OsRbohB-FLAG by PEG-mediated transformation. GFP was expressed as a negative control. After incubation for 14–16 h, rice protoplasts were collected for protein extraction using extraction buffer (50 mM Tris-HCl, pH 7.4, 150 mM NaCl, 1 mM EDTA, 1% Triton X-100) supplemented with protease inhibitor cocktail (CWBIO, Beijing, China).

Total protein extracts were incubated with 25 μl anti-FLAG M2 affinity beads (Sigma-Aldrich, St. Louis, MO, USA) at 4 °C for 4 h with gentle rotation. The beads were washed thoroughly with ice-cold PBS containing protease inhibitors. Bound proteins were eluted by heating the beads at 95 °C for 5 min in SDS-PAGE loading buffer. Immunoprecipitated proteins were resolved by SDS-PAGE and subjected to immunoblotting with HRP-conjugated anti-FLAG antibody (Sigma-Aldrich, St. Louis, MO, USA; 1:5,000) and anti-phosphoserine/threonine antibody (Millipore, Burlington, MA, USA; 1:5,000). Input samples were also detected by immunoblotting with an anti-Actin antibody (CWBIO, Beijing, China; 1:5,000). Signals were detected using an enhanced chemiluminescence detection system.

### Protein structure modelling

The structural model of OsRbohB was generated by AlphaFold 3^48^. The human NOX5 Ca^2+^-binding structure (PDB: 6SZ5) was retrieved from the Protein Data Bank and used for structural comparison^49^. The OsRbohB model and the human NOX5 Ca^2+^-binding structure were aligned using the Matchmaker tool in UCSF ChimeraX^50^, focusing on the EF-hand Ca^2+^-binding loop and adjacent α-helices.

### Microscale thermophoresis assay

Microscale thermophoresis (MST) assays were performed as described previously with minor modifications^51^. DNA fragments encoding OsRbohB^1-3^^55^, OsRbohB^1-3^^55(T288D)^, OsRbohI^1-383^, and OsRbohI^1-383(S317D)^ were amplified and fused in frame downstream of the GFP coding sequence amplified from pRTV-GFP^52^. The resulting GFP-OsRboh fusion fragments were cloned into pGEX-4T-1 to express GST-tagged GFP-fusion proteins. The constructs were transformed into *Escherichia coli* BL21(DE3) cells for recombinant protein expression, and the fusion proteins were purified for MST analysis.

Purified GST-GFP-OsRboh proteins were diluted in MST buffer (50 mM Tris-HCl, pH 7.5, 150 mM NaCl and 0.05% Tween 20). A constant concentration of purified protein was mixed with serial dilutions of CaCl_2_ and incubated overnight at 4 °C before measurement. Binding measurements were performed on a Monolith NT.115 instrument (NanoTemper Technologies, Munich, Germany) according to the manufacturer’s protocol. Dissociation constants were calculated by fitting the normalized fluorescence changes to a one-site binding model.

### ROS burst assay

ROS production was measured using a luminol-based chemiluminescence assay adapted from established protocols^53^. The coding sequences encoding OsRboh variants were cloned into the pGD vector to express OsRboh-GFP fusion proteins^54^. *N. benthamiana* leaves were infiltrated with *Agrobacterium tumefaciens* strains carrying *OsRboh GFP* variants. Leaf discs (3 mm diameter) were collected at 2 days after agroinfiltration and floated overnight in 100 µl sterile H_2_O per well of a 96-well plate at room temperature in the dark. Rice leaf discs were prepared and equilibrated using the same procedure. For elicitation, 100 μl of reaction solution containing 100 μM luminol (FUJIFILM Wako Pure Chemical Corporation, Osaka, Japan), 40 μg ml^-1^ horseradish peroxidase (HRP; Coolaber, Beijing, China) and either 20 μg ml^-1^ chitin (Sigma-Aldrich, St. Louis, MO, USA; Cat. no. C9752), 2 μM flg22 (Sangon Biotech, Shanghai, China) or mock solution was added to each well, giving a final volume of 200 μl. Chemiluminescence was recorded immediately at 1 min intervals for 40 min using a Spark 20M multimode microplate reader (Tecan, Männedorf, Switzerland).

### Cell culture and ROS assay in HEK293T cells

ROS production in HEK293T cells was measured using a chemiluminescence assay adapted from Han *et al.* (2019)^25^. Briefly, HEK293T cells were maintained in Dulbecco’s modified Eagle’s medium (DMEM; Biosharp, Beijing, China) supplemented with 10% fetal bovine serum (FBS; Biological Industries, Beit Haemek, Israel) and 1% penicillin–streptomycin at 37 °C in a 5% CO_2_ atmosphere. The coding sequence of *OsRbohB* and the fragment encoding OsCPK17-VK (residues 1-400) were cloned into pEF expression vectors. For ROS assays, HEK293T cells were seeded in 96 well plates and transfected with the indicated plasmids using polyethylenimine (PEI; XANCON, Beijing, China) according to standard protocols.

At 48 h after transfection, cells were washed with calcium- and magnesium-free Hanks’ balanced salt solution (HBSS) without phenol red (Shanghai Yuanye Bio-Technology Co., Ltd., Shanghai, China) and incubated in 100 μl of ROS detection buffer containing 250 μM L-012 (FUJIFILM Wako Pure Chemical Corporation, Osaka, Japan) and 6 U ml^-1^ horseradish peroxidase (HRP; Coolaber, Beijing, China). Basal luminescence was recorded for 10 min using a FlexStation 3 microplate reader (Molecular Devices, San Jose, CA, USA). Subsequently, 50 μl of the ROS detection buffer supplemented with 3 mM CaCl_2_ and 3 μM ionomycin (Sigma-Aldrich, St. Louis, MO, USA) was automatically injected into each well, and luminescence was continuously recorded for an additional 15 min.

### *M. oryzae* inoculation and fungal biomass quantification

*M. oryzae* inoculation assay was performed as described previously with minor modifications^55^. The *M. oryzae* isolate RB22 was cultured on oat/V8 medium plates at 28 °C in darkness for 1 week and then exposed to continuous fluorescent light at room temperature for an additional week to induce sporulation. Conidia were harvested and resuspended in sterile water containing 0.025% (v/v) Tween 20 to a final concentration of 5×10^5^ conidia ml^-1^. Three-week-old rice seedlings were evenly sprayed with conidial suspension, kept in darkness under high-humidity conditions for 24 h, and then transferred to normal growth conditions.

Disease symptoms were photographed at 7 d after inoculation, and lesion areas were quantified from digital images using Adobe Photoshop (Adobe, San Jose, CA, USA). For fungal biomass quantification, genomic DNA was extracted from infected leaves. Relative fungal biomass was determined by DNA-based quantitative PCR using the *M. oryzae* transposable element *MoPot2* and rice *UBIQUITIN* as the fungal and host genomic markers, respectively.

### *U. virens* inoculation assay

*U. virens* inoculation assay was performed as described previously with minor modifications^56^. The *U. virens* strain U14Kc002 was cultured on potato sucrose agar (PSA) plates at 28 °C for 10 d and then transferred into potato sucrose (PS) liquid medium for 7 d at 28 °C with shaking at 150 rpm. The culture containing hyphae and conidia was homogenized and diluted in PS medium to a final concentration of 1 × 10^6^ spores ml^-1^. The resulting suspension was injected into rice panicles at 5-7 d before heading. At least 10 panicles were inoculated for each genotype. False smut balls formed on inoculated rice panicles were counted at 4-6 weeks after inoculation.

### *X. oryzae* pv. *oryzicola* inoculation assay

*X. oryzae* pv. *oryzicola* strain RS105 was used for bacterial leaf streak inoculation assays as described previously with minor modifications^57^. RS105 cells were first cultured on nutrient agar plates supplemented with 25 μg ml^-1^ rifampicin at 28 °C for 2 d, and then grown to exponential phase in rifampicin-containing (25 μg ml^-1^) nutrient broth at 28 °C with shaking for 16 h. Bacterial cells were collected by centrifugation, washed with sterile water and resuspended in 10 mM MgCl_2_ to an OD_600_ of 0.5. Fully expanded leaves of six-week-old rice plants were infiltrated with bacterial suspension using a needleless syringe. Disease symptoms were recorded at 8 d after inoculation, and lesion lengths were measured to evaluate bacterial leaf streak resistance.

### Defense marker gene expression analysis

Defense marker gene expression was analysed by quantitative reverse transcription PCR (RT-qPCR) as described previously^46^. Seven-day-old rice seedlings grown aseptically on half-strength Murashige and Skoog (1/2 MS) medium were treated with 10 μg ml^-1^ chitin, 1 µM flg22 or mock solution as indicated. Leaf tissues were collected at the indicated time points, immediately frozen in liquid nitrogen and stored at −80 °C until RNA extraction. Total RNA was extracted using an Ultrapure RNA isolation kit (Mei5bio, Beijing, China) according to the manufacturer’s instructions. First-strand cDNA was synthesized from DNase-treated RNA using a reverse transcription kit (Mei5bio, Beijing, China). RT-qPCR was performed using gene-specific primers and SYBR Green qPCR mix (Mei5bio, Beijing, China). Relative transcript levels were calculated using the 2^−ΔΔCt^ method, with *OsActin1* used as the internal reference gene.

### Prime editing and genotyping

Prime-edited rice plants were generated using a modified PE-P3 prime-editing system^58^. The prime editor contains an N-terminal Moloney murine leukaemia virus reverse transcriptase fused to the H840A nickase variant of *Streptococcus pyogenes* Cas9. Prime-editing guide RNAs were designed to restore ancestral acidic residues in two rice Rboh proteins.

For OsRbohB, the codon encoding Thr288 in LOC_Os01g25820.1 was converted from ACA to GAT to generate the *OsRbohB^T288D^* allele. The pegRNA protospacer was 5’-TCGATGTATCCCAAGTTTGT-3’, and the non-edited-strand nicking sgRNA protospacer was 5’-AAACAAACTTTCCAAGATCA-3’. The corresponding wild-type target-region sequence was 5’-AAGAGCTTGACCCTACAAACTTGGG-3’, and the programmed edited sequence was 5’-AAGAGCTTGACCCTGATAACTTGGG-3’.

For OsRbohI, the codon encoding Ser317 in LOC_Os11g33120.1 was converted from AGC to GAC to generate the *OsRbohI^S317D^* allele. The pegRNA protospacer was 5’-TCGATGTAGCCGAGGTTGCT-3’, and the non-edited-strand nicking sgRNA protospacer was 5’-GTCGGCCAACAAGCTGTCCA-3’. The wild-type target-region sequence was 5’-TCGACCCCAGCAACCTCGGC-3’, and the programmed edited sequence was 5’-TCGATCCCGACAACCTCGGC-3’. This design introduced the intended Ser317Asp substitution together with an additional synonymous substitution.

The pegRNA and nicking sgRNA expression cassettes were amplified using primers containing *Bsa* I sites and assembled into the modified PE-P3 backbone after *Bsa* I digestion. The guide, reverse-transcription template and primer-binding-site regions of each construct were verified by Sanger sequencing. The resultant binary vectors were introduced into *A. tumefaciens* strain EHA105 and transformed into embryogenic calli induced from mature seeds of ZH11. Hygromycin-resistant calli were regenerated to obtain T0 plants.

Transgenic plants carrying the prime-editing construct were identified by PCR using vector-specific primers, and genomic regions spanning the target sites in *OsRbohB* and *OsRbohI* were amplified using locus-specific primers (Table S4). PCR amplicons were subjected to Sanger sequencing to identify plants carrying precise programmed edits. Edited plants were self-pollinated and independent homozygous progeny lines carrying the desired *OsRbohB^T288D^* or *OsRbohI^S317D^* allele were selected for subsequent analyses.

### OsRbohB orthologue identification and Thr288 residue-state analysis

To identify OsRbohB orthologues in Poales, the nine Rboh protein sequences from *O. sativa* were used as queries in BLASTP searches against the predicted proteomes of 16 additional Poales species. The searched proteomes and identified OsRbohB orthologues, including the *O. sativa* reference sequence, are listed in Table S7. For each proteome, a local BLAST database was constructed using the predicted protein sequences. BLASTP searches were performed with an E-value threshold of 1 × 10^-5^, a maximum of 100,000 target sequences per query, and tabular output format^59^. For each BLASTP hit, global pairwise amino-acid identity between the hit protein and each of the nine rice Rboh proteins was calculated using the Biostrings package in R. A protein was defined as an OsRbohB orthologue if it showed the highest sequence identity to OsRbohB among all nine rice Rboh proteins, with the threshold of this identity set to exceed 50%. The amino-acid residue corresponding to OsRbohB Thr288 was extracted from the alignment of each identified orthologue. Species in which no protein met these criteria were considered to lack an identifiable OsRbohB orthologue.

To investigate the residues at OsRbohB Thr288-equivalent site in cultivated rice, sequence variation and subpopulation information for 4,419 *O. sativa* accessions were obtained from RiceVarMap2^60^. All variants overlapping the codon encoding Thr288 in LOC_Os01g25820.1 were systematically screened across the full dataset and used to infer the encoded amino-acid residue in each accession. Residues were summarized by rice subpopulation according to the RiceVarMap2 classification.

### Field trial under contrasting fertilization regimes

To evaluate the effect of fertilization regime on agronomic traits of *OsRbohB^T288D^* prime-edited lines, we conducted a field trial in a previously uncultivated field at Nanbin Farm, Sanya, Hainan, China. The field was divided into eight 50 m^2^ subplots with two fertilization regimes and four subplots per regime according to the layout shown in Figure S11. Soil bunds approximately 15 cm high were constructed around each subplot and covered with plastic film to prevent the lateral movement of water and nutrients between adjacent subplots. Rice seedlings were grown for 28 d before transplanting and were transplanted into all subplots at one seedling per hill, with 25 cm between rows and 14 cm between hills.

For the fertilized regime, four subplots received a potassium-sulfate-type compound fertilizer, 15-15-15 N-P_2_O_5_-K_2_O (Kingenta Ecological Engineering Group Co., Ltd., China), at 1 kg per 50 m^2^ subplot per application at 4, 14 and 32 d after transplanting. The remaining four subplots received no compound fertilizer throughout the experiment. Except for the fertilization regime, all field management practices, including land preparation, transplanting, weed control, fungicide and insecticide application, and protective netting, were applied uniformly across all subplots. Plants were grown to maturity under field conditions. For each genotype under each fertilization regime, plants or panicles were randomly sampled from the four corresponding subplots to measure seed-setting rate, filled grains per panicle, hundred-grain weight and grain weight per plant.

### Agronomic trait evaluation

Agronomic traits were evaluated as described previously^61^. Plant height was measured from the soil surface to the tip of the uppermost panicle at maturity. The number of effective tillers per plant was recorded as the number of tillers bearing fully developed panicles. Seed-setting rate was determined using panicles sampled from comparable positions on individual plants and calculated as the ratio of filled grains to total grains per panicle. For single-plant grain weight, all filled grains from individual plants were harvested and weighed. Random subsets of 100 mature filled grains were selected for the hundred-grain weight.

## QUANTIFICATION AND STATISTICAL ANALYSIS

Statistical analyses were performed using GraphPad Prism version 8.0 (GraphPad Software, San Diego, CA, USA). Data are presented as mean ± s.e.m. unless otherwise indicated. For comparisons among multiple groups, ordinary one-way analysis of variance (ANOVA) followed by Tukey’s multiple comparisons test was used to compare the means among individual groups. For pairwise comparisons, two-tailed Student’s *t*-tests were performed. Differences were considered statistically significant at *P* < 0.05.

---

**Figure S1.**
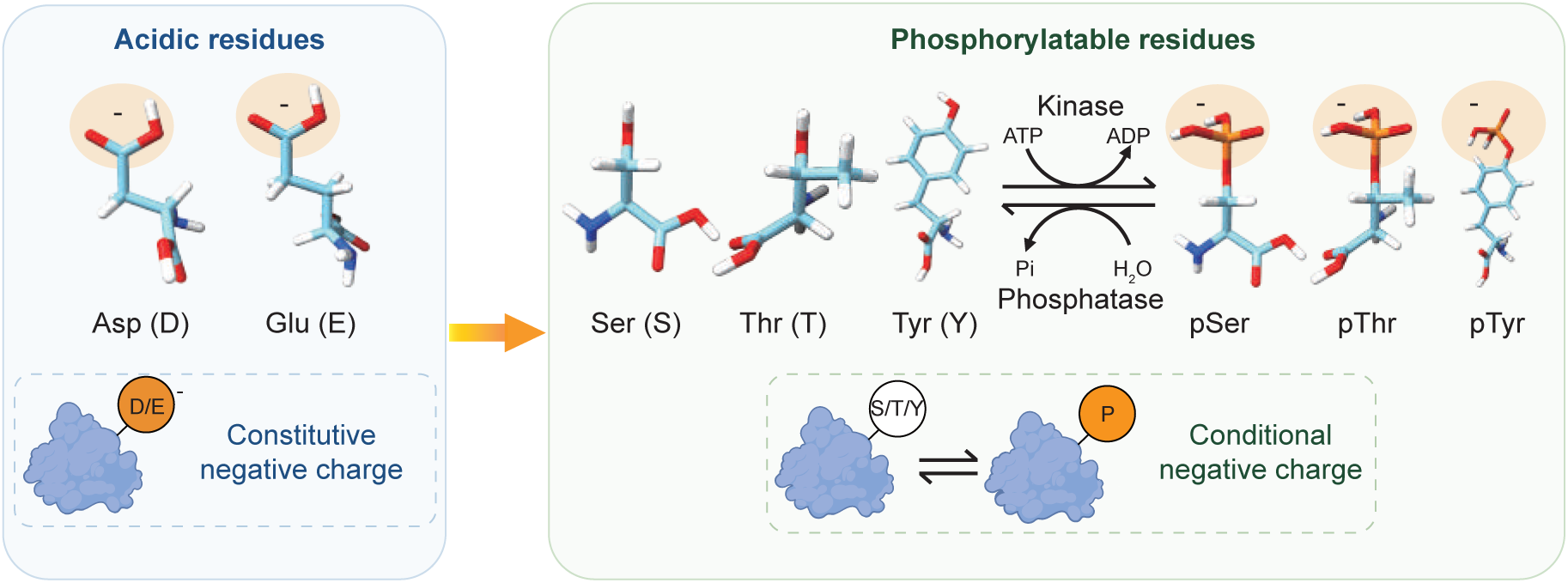
Evolutionarily acquired phosphorylation sites create reversible negative charge states. Conceptual model illustrating that a subset of phosphorylatable residues, Ser (S), Thr (T) or Tyr (Y), can arise through substitutions from ancestral acidic residues Asp (D) or Glu (E). Acidic residues usually provide a stable negative charge under physiological conditions, whereas phosphorylation of Ser, Thr or Tyr by protein kinases reversibly restores this negative charge, which phosphatases can remove, creating a mechanism for signal-dependent regulation via evolutionarily acquired phosphorylation sites.

**Figure S2.**
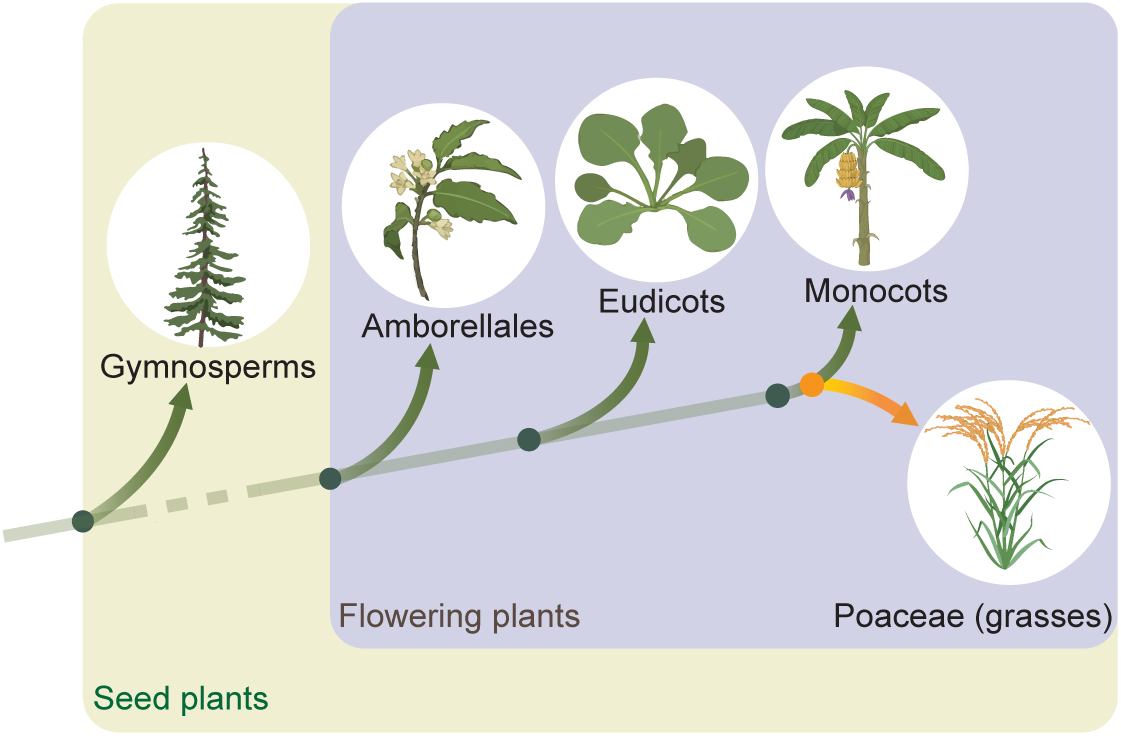
Phylogenetic context of Poaceae within seed plants. Simplified phylogenetic framework showing the placement of Poaceae within major seed plant lineages. Gymnosperms are shown as the sister lineage to flowering plants, and flowering plants are represented by Amborellales, Eudicots and Monocots. Poaceae, the grass family that includes rice, is highlighted within monocots. Illustrations depict representative members of each lineage. The schematic is not drawn to scale.

**Figure S3.**
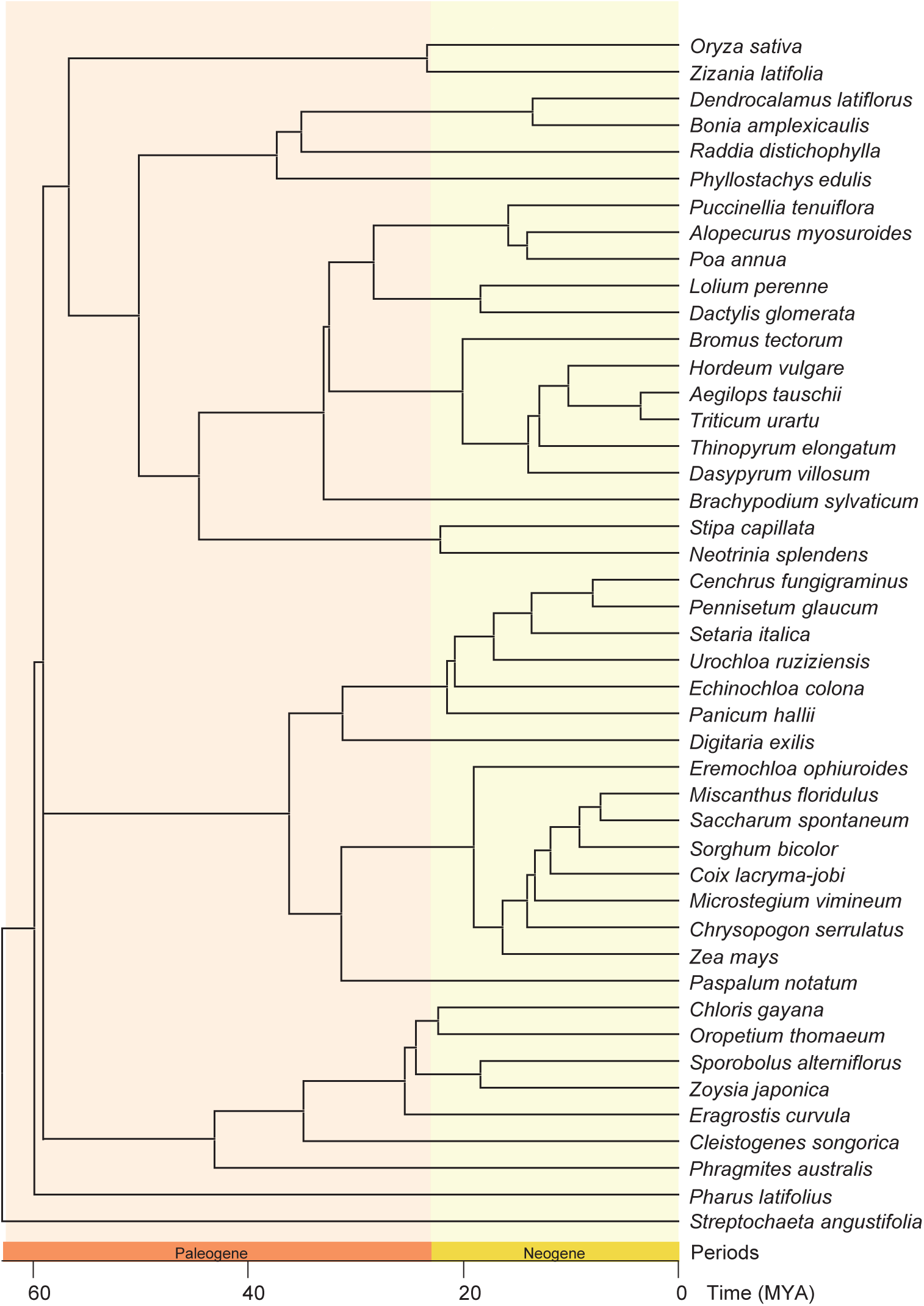
Expanded phylogenetic tree of 45 Poaceae species used for whole-genome alignment. Expanded phylogenetic tree of the 45 Poaceae species used for whole-genome alignment and phyloP-based prioritization of candidate rice EAPSs. The phylogenetic framework and divergence times were obtained from the TimeTree database. Each terminal branch represents an individual species. The timescale is shown in millions of years ago (MYA), and the bottom color bar denotes geological periods.

**Figure S4.**
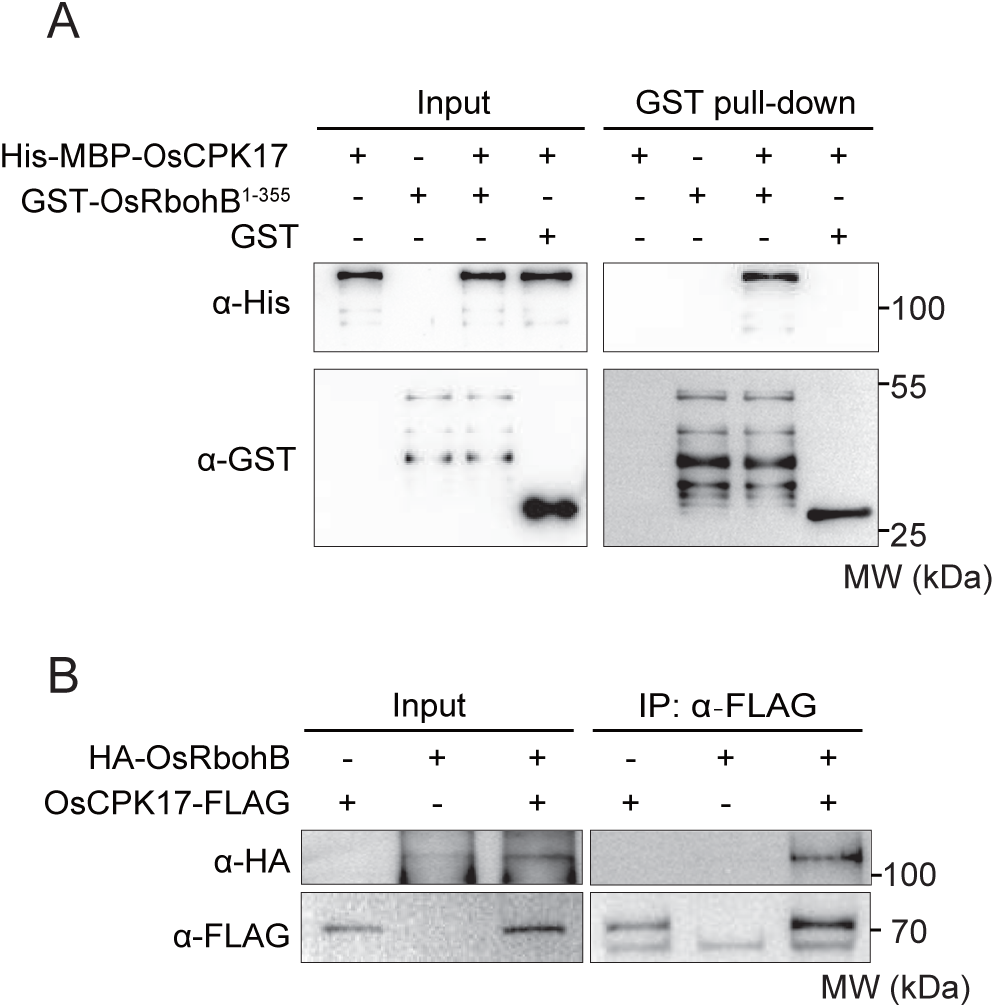
OsCPK17 interacts with OsRbohB *in vitro* and in rice protoplasts. (A) GST pull-down assay showing the interaction between the N-terminal cytosolic region of OsRbohB and OsCPK17. GST-OsRbohB^1-3^^55^ or GST was incubated with His-MBP-OsCPK17 in the presence of GST beads. Bead-bound proteins were detected by immunoblotting with anti-His and anti-GST antibodies. (B) Co-immunoprecipitation assay showing the interaction between OsRbohB and OsCPK17 in rice protoplasts. HA-OsRbohB and OsCPK17-FLAG were individually expressed or co-expressed in rice protoplasts. Protein extracts were immunoprecipitated with anti-FLAG affinity beads and analysed by immunoblotting with anti-HA and anti-FLAG antibodies. MW, molecular weight.

**Figure S5.**
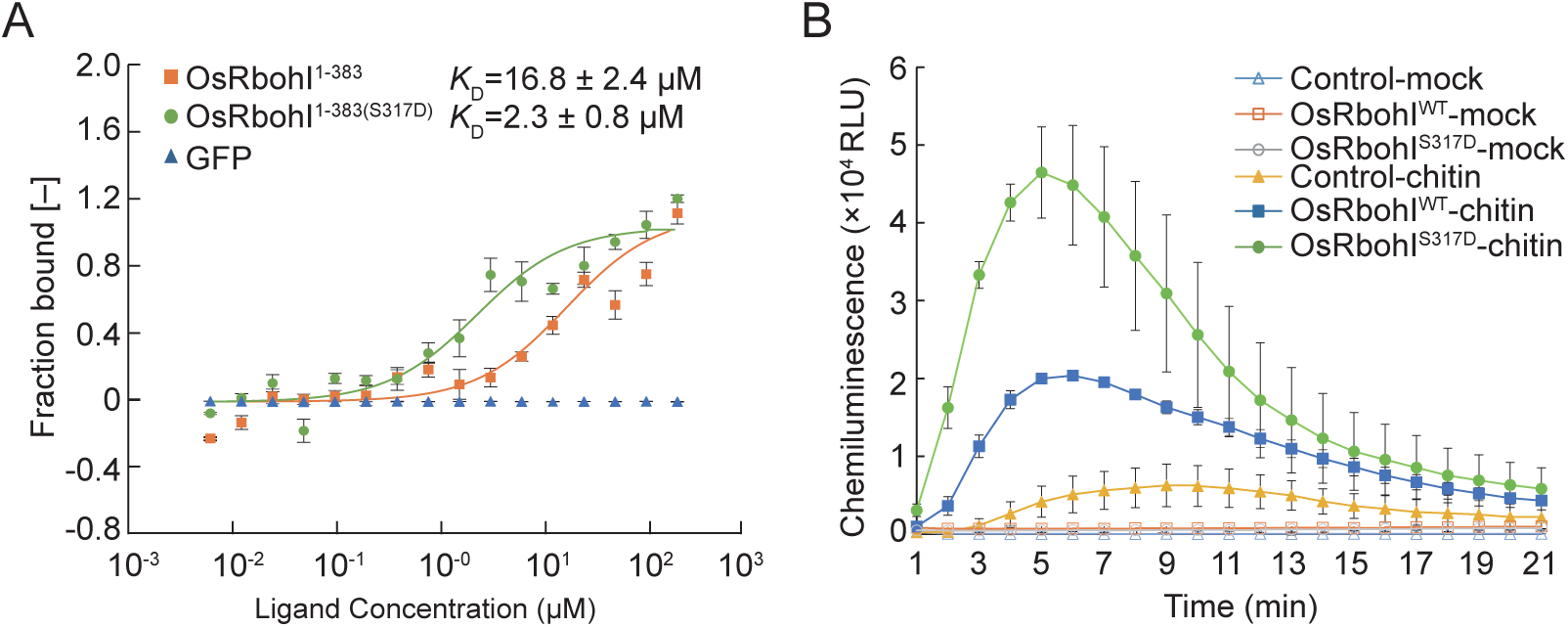
Restoration of the ancestral residue at OsRbohI Ser317 enhances Ca^2+^ binding and ROS production. (A) Microscale thermophoresis analysis of Ca^2+^ binding by purified GFP, OsRbohI^1-383^ and OsRbohI^1-383(S317D)^ proteins. Curves show one-site binding fits; apparent dissociation constants (*K*_D_) are indicated. Data are mean ± s.e.m.; n = 3. (B) Chitin-induced ROS burst in *N. benthamiana* leaves infiltrated with an *Agrobacterium*-only control or transiently expressing OsRbohI^WT^ or OsRbohI^S317D^. ROS production was measured by luminol-based chemiluminescence after mock or chitin treatment. Data are mean ± s.e.m.; n = 8.

**Figure S6.**
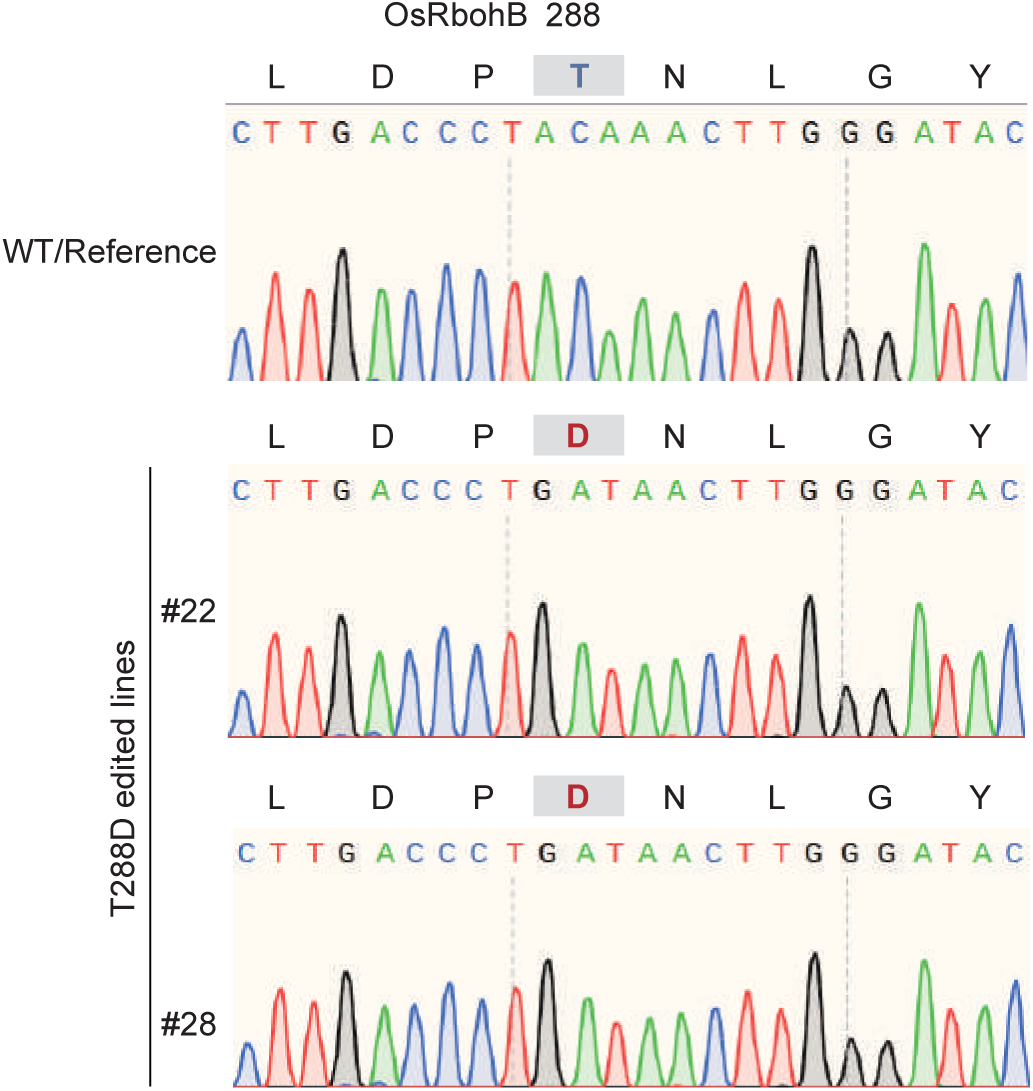
Genotyping of prime-edited rice *OsRbohB^T288D^* mutant lines by Sanger sequencing. DNA sequencing chromatograms of *OsRbohB^T288D^* prime-edited rice mutant lines at target site of *OsRbohB*.

**Figure S7.**
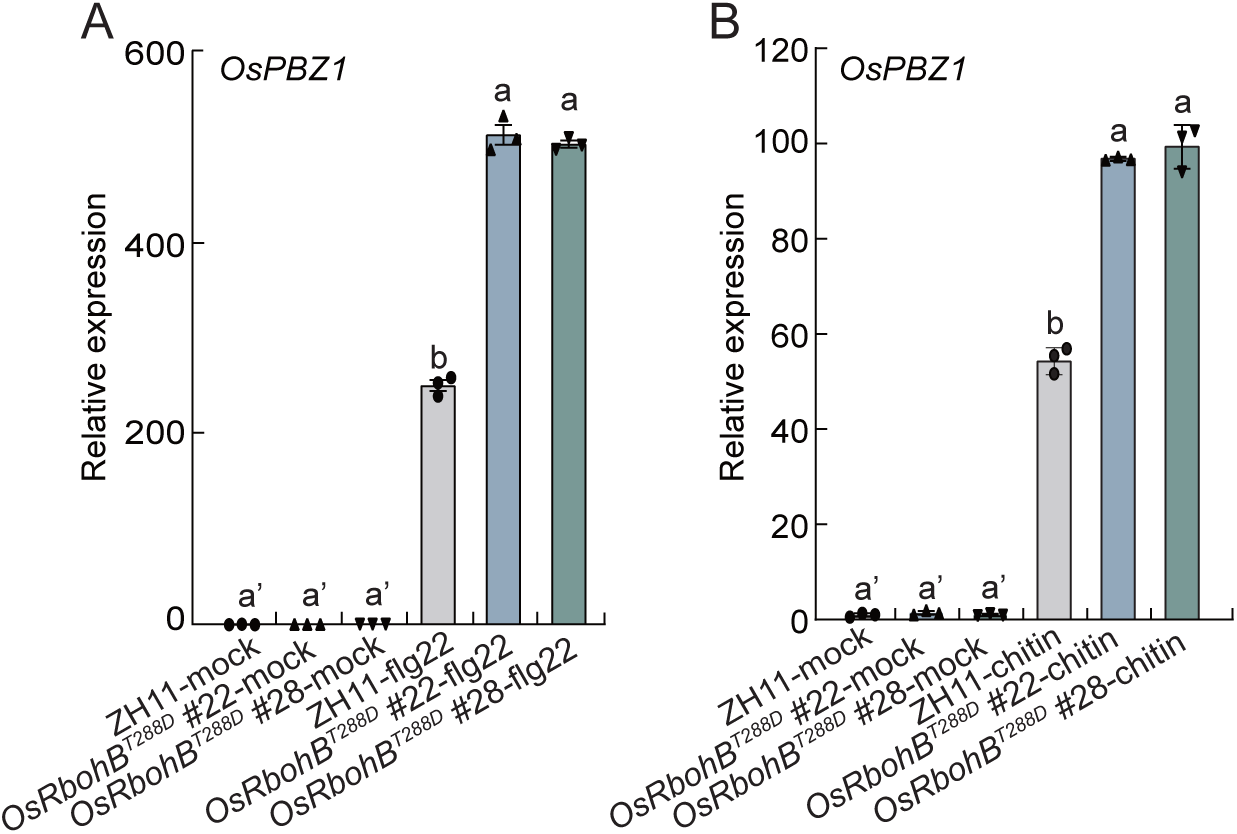
Prime-edited *OsRbohB^T288D^* rice lines show enhanced PAMP-induced *OsPBZ1* expression. (A and B) Relative expression levels of *OsPBZ1* in wild-type ZH11 and two independent *OsRbohB^T288D^* prime-edited lines after flg22 (A) or chitin (B) treatment. Transcript levels were normalized to *OsActin1*. Data are mean ± s.e.m.; n = 3. Different letters indicate statistically significant differences among the indicated groups.

**Figure S8.**
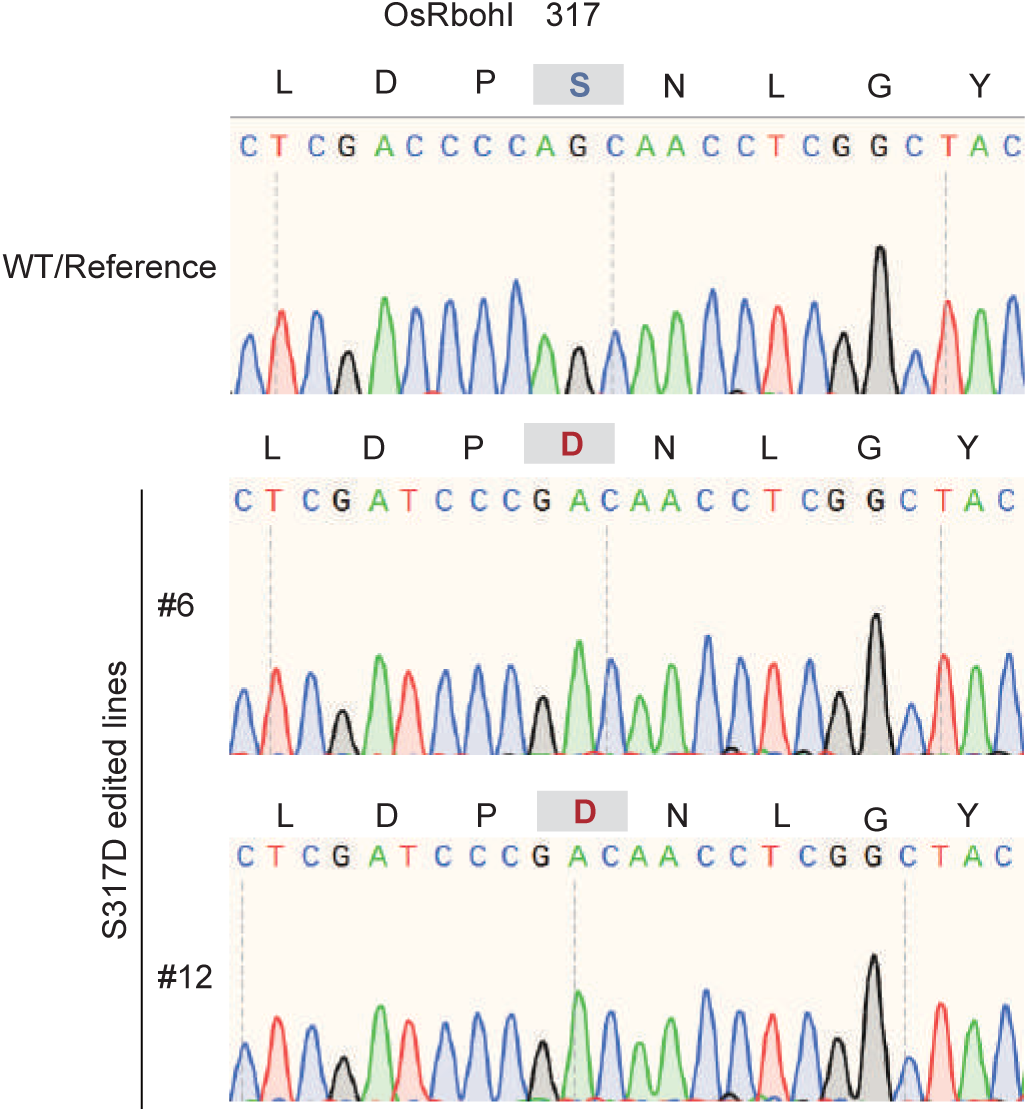
Genotyping of prime-edited rice *OsRbohI^S317D^* mutant lines by Sanger sequencing. DNA sequencing chromatograms of *OsRbohI^S317D^* prime-edited rice mutant lines at target site of *OsRbohI*.

**Figure S9.**
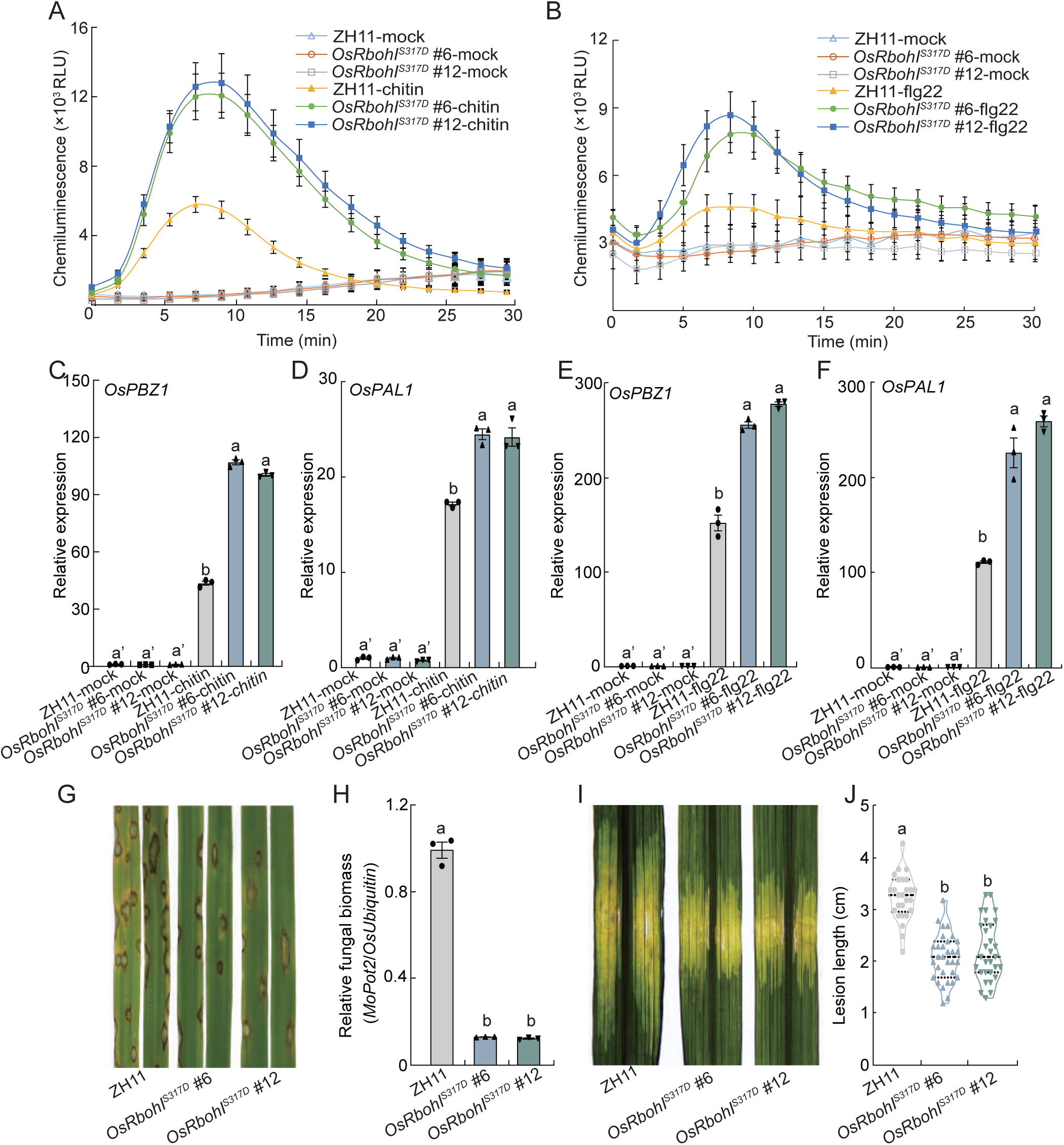
Ancestral-state restoration at OsRbohI Ser317 enhances PAMP-triggered immunity and disease resistance. (A and B) Chitin-induced (A) and flg22-induced (B) ROS bursts in leaves of wild-type ZH11 and two independent *OsRbohI^S317D^* prime-edited lines. ROS production was measured by luminol-based chemiluminescence. Data are mean ± s.e.m.; n = 16. (C-F) Relative expression levels of *OsPBZ1* (C and E) and *OsPAL1* (E and F) in ZH11 and *OsRbohI^S317D^* plants after chitin (C and D) or flg22 (E and F) treatment. Transcript levels were normalized to *OsActin1* expression. Data are mean ± s.e.m.; n = 3. Different letters indicate statistically significant differences among the indicated groups. (G and H) Representative rice blast lesions (G) and relative fungal biomass (H) in ZH11 and *OsRbohI^S317D^* lines after inoculation with *M. oryzae* isolate RB22. Fungal biomass was quantified by DNA-based qPCR. Data are mean ± s.e.m.; n = 3. Different letters indicate statistically significant differences among the indicated groups. (I and J) Representative bacterial leaf streak symptoms (I) and lesion-length quantification (J) in ZH11 and two independent *OsRbohI^S317D^* lines after inoculation with *X. oryzae* pv. *oryzicola* strain RS105. Horizontal lines indicate the median and quartiles. Different letters indicate statistically significant differences among genotypes.

**Figure S10.**
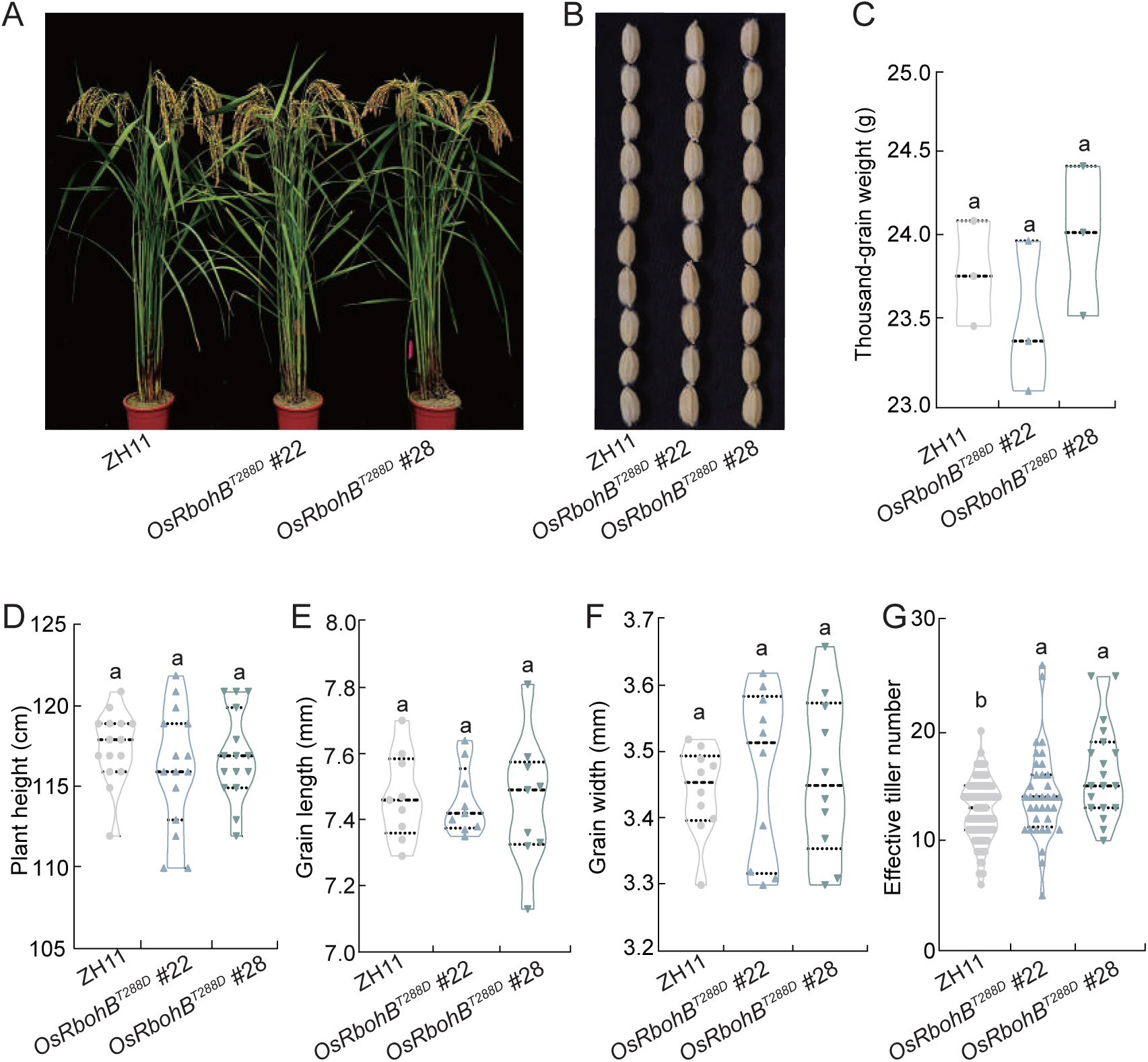
Major agronomic traits of prime-edited *OsRbohB^T288D^* rice lines under fertilized cultivation. (A and B) Representative images of mature plants (A) and harvested grains (B) from wild-type ZH11 and two independent *OsRbohB^T288D^* prime-edited lines grown under fertilized field conditions. (C-G) Quantification of thousand-grain weight (C), plant height (D), grain length (E), grain width (F) and effective tiller number (G) in ZH11 and *OsRbohB^T288D^* lines. Violin plots show the distribution of individual values; horizontal lines indicate the median and quartiles. Different letters indicate statistically significant differences among genotypes.

**Figure S11.**
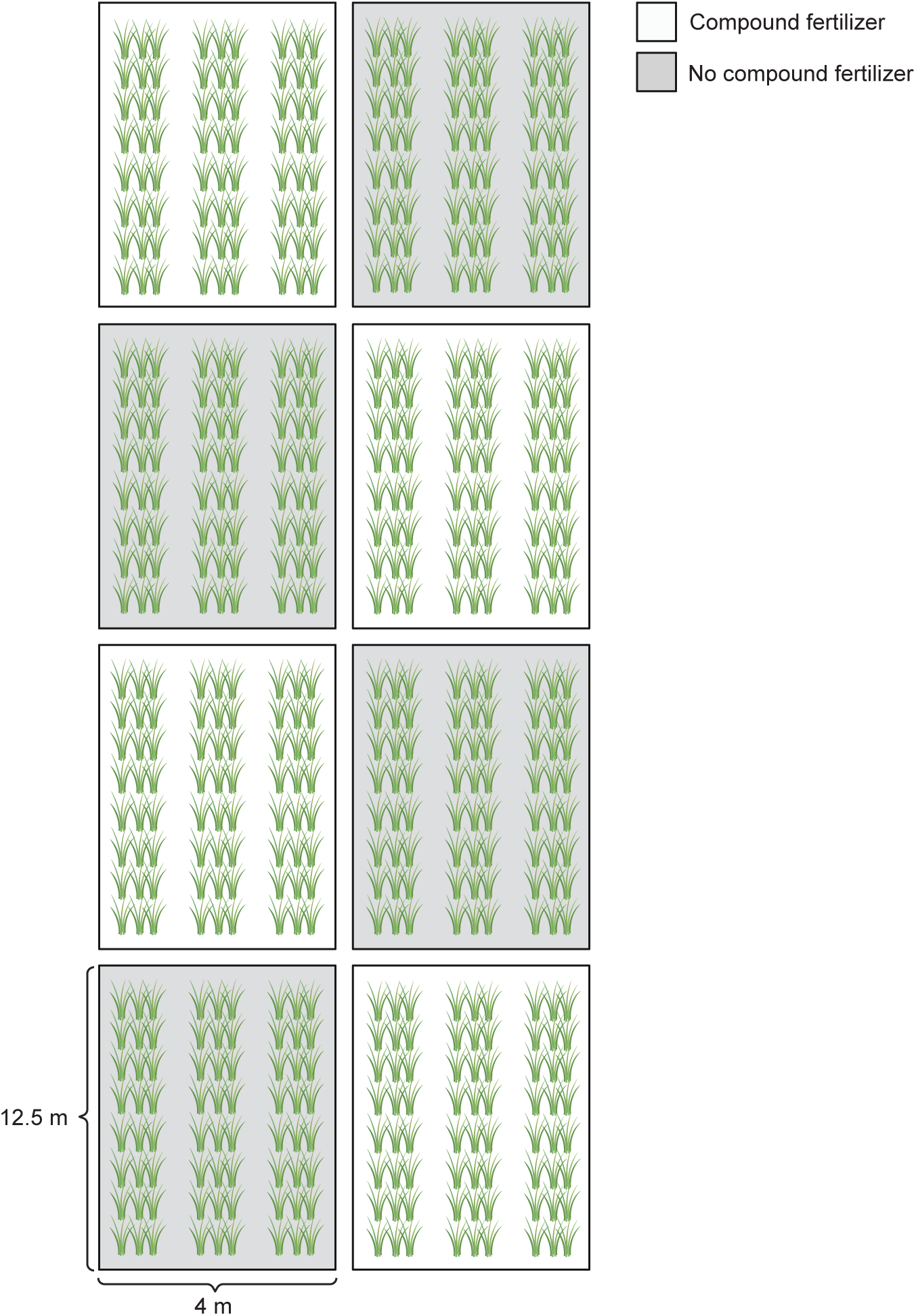
Field layout of the fertilization trial. The field was divided into eight subplots, each measuring 4 m × 12.5 m (50 m^2^). Within each subplot, the ZH11, *OsRbohB^T288D^* #22 and *OsRbohB^T288D^* #28 rice lines were grown in separate rows. White boxes indicate subplots that received compound fertilizer, and grey boxes indicate subplots that received no compound fertilizer. The fertilized subplots received 1 kg of compound fertilizer per subplot per application at 4, 14 and 32 d after transplanting.

**Figure S12.**
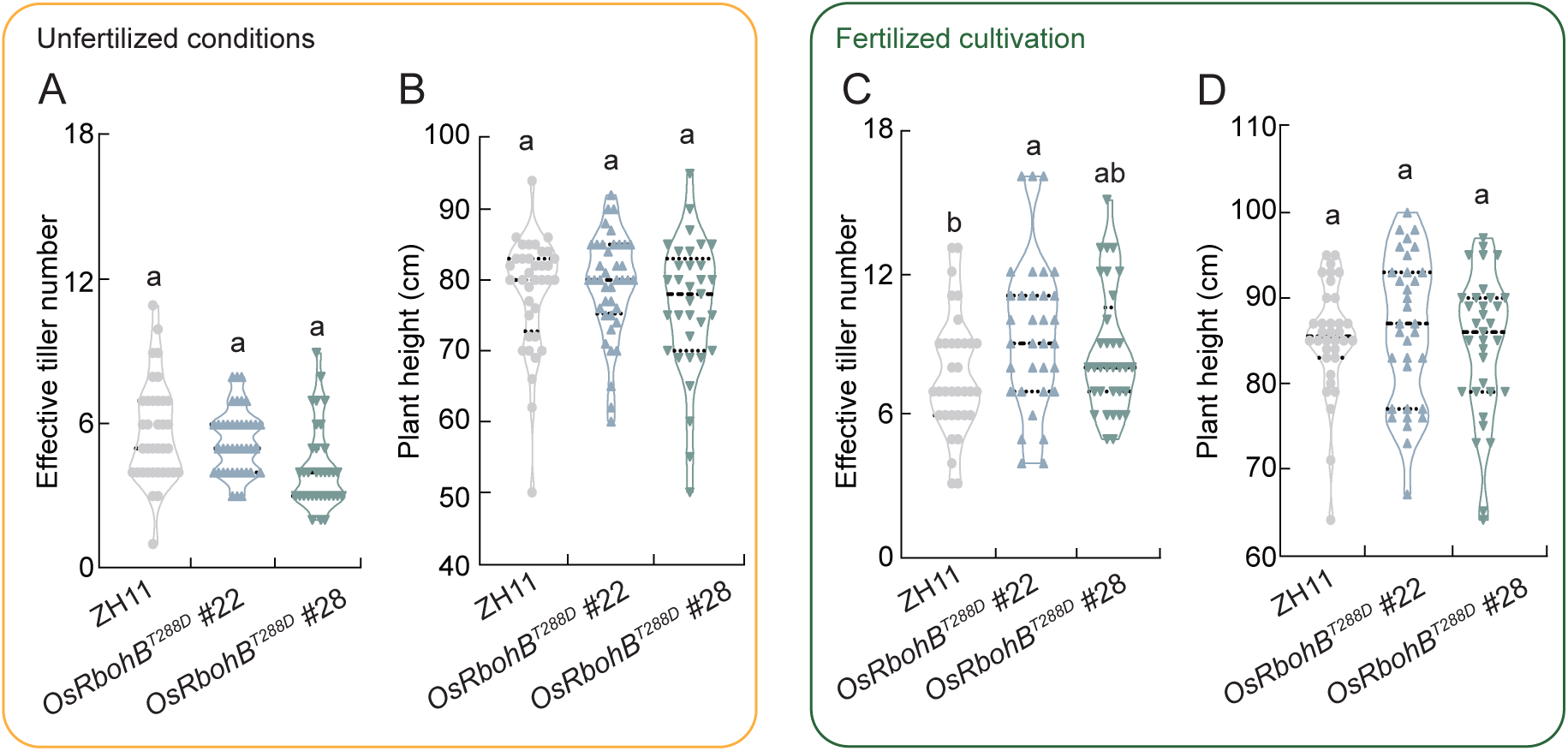
Effective tiller number and plant height of prime-edited *OsRbohB^T288D^* rice lines under contrasting nutrient conditions. (A and B) Effective tiller number (A) and plant height (B) of wild-type ZH11 and two independent *OsRbohB^T288D^* prime-edited lines in the paddy field without fertilizer application. (C and D) Effective tiller number (C) and plant height (D) of ZH11 and *OsRbohB^T288D^* lines under fertilized cultivation. Violin plots show the distribution of individual values; horizontal lines indicate the median and quartiles. Different letters indicate statistically significant differences among genotypes.

